# Mu-Transcranial Alternating Current Stimulation Induces Phasic Entrainment and Plastic Facilitation of Corticospinal Excitability

**DOI:** 10.1101/2022.10.17.512611

**Authors:** Asher Geffen, Nicholas Bland, Martin V Sale

## Abstract

Transcranial alternating current stimulation (tACS) has been proposed to modulate neural activity through two primary mechanisms: entrainment and neuroplasticity. The current study aimed to probe both of these mechanisms in the context of the sensorimotor µ-rhythm using transcranial magnetic stimulation (TMS) and electroencephalography (EEG) to assess entrainment of corticospinal excitability (CSE) during stimulation (i.e., online) and immediately following stimulation, as well as neuroplastic aftereffects on CSE and µ EEG power. Thirteen participants received 3 sessions of stimulation. Each session consisted of 90 trials of µ-tACS tailored to each participant’s individual µ frequency (IMF), with each trial consisting of 16 seconds of tACS followed by 8 seconds of rest (for a total of 24 minutes of tACS and 12 minutes of rest per session). Motor evoked potentials (MEPs) were acquired at the start and end of the session (n = 41) and additional MEPs were acquired across the different phases of tACS at 3 epochs within each tACS trial (n = 90 for each epoch): early online, late online, and offline echo. Resting EEG activity was recorded at the start, end, and throughout the tACS session. The data were then pooled across the three sessions for each participant to maximise the MEP sample size per participant. We present preliminary evidence of CSE entrainment persisting immediately beyond tACS and have also replicated the plastic CSE facilitation observed in previous µ-tACS studies, thus supporting both entrainment and neuroplasticity as mechanisms by which tACS can modulate neural activity.

**Graphical Abstract:** Thirteen participants underwent 3 sessions of stimulation where they received 90 trials of mu-tACS (270 trials across the 3 sessions), with each trial consisting of 16 seconds of tACS (2mA at the participants individual mu frequency) followed by 8 seconds of rest. Motor evoked potentials (MEPs) were acquired at the start and end of the session (n = 41) and additional MEPs were acquired across the different phases of tACS at 3 epochs within each tACS trial (n = 90 for each epoch): early online, late online, and offline echo. We present preliminary evidence supporting entrainment of MEP amplitudes to tACS phase online to and immediately following stimulation and have also replicated the neuroplastic CSE facilitation observed in previous µ-tACS studies.

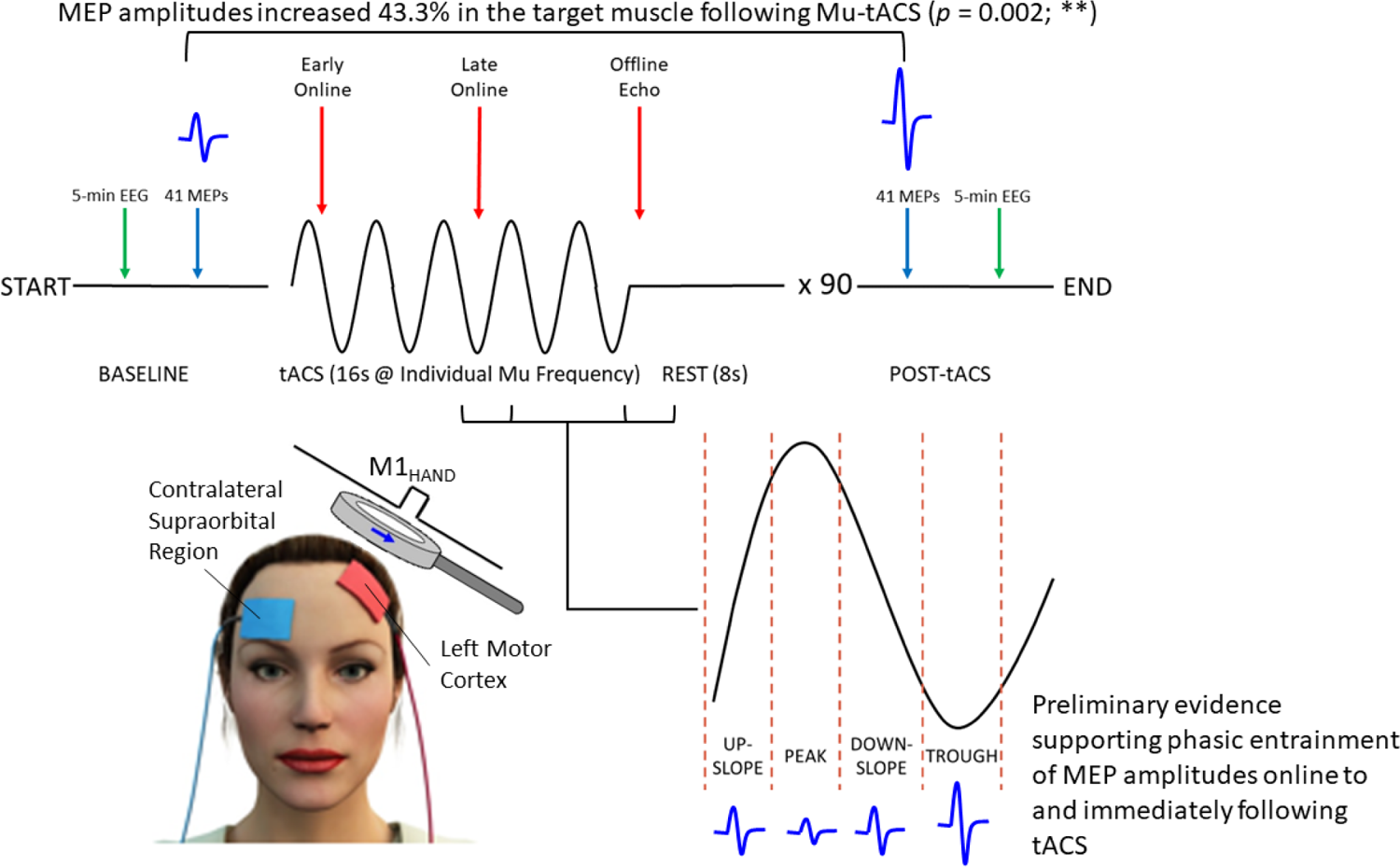

## 1. Introduction

Over the last two decades, researchers have used a type of non-invasive brain stimulation called transcranial alternating current stimulation (tACS) to experimentally modulate various aspects of behaviour and cognition (e.g., attention, memory, perception, and motor processes) by matching the frequency of the applied alternating current with the frequency of a given neural oscillation known to be associated with a particular behavioural/cognitive task (for reviews see Antal and Paulus, 2013; Bland and Sale, 2019; Herrmann et al., 2016; Vosskuhl et al., 2018). However, despite promising behavioural/cognitive effects, a thorough understanding of the underlying neurophysiological mechanisms is still required if this technique is to be used across a wide range of clinical settings, such as in psychiatry (e.g., Alexander et al., 2019) and neurorehabilitation (e.g., Del Felice et al., 2019).

At the neuronal level, tACS induces oscillatory shifts in the membrane potentials of pyramidal cells between states of depolarisation and hyperpolarisation (Vöröslakos et al., 2018). Because the stimulation is sub-threshold (i.e., the voltage changes induced in the membrane potentials of the affected neurons are not sufficient to directly cause those neurons to depolarise; Elyamany et al., 2021; Vosskuhl et al., 2018), tACS does not necessarily alter the firing rate of action potentials, but rather probabilistically affects their spike timing in a manner that is specific to the frequency and phase of the applied stimulation (Reato et al., 2010).

At the network level, there is growing evidence from animal (e.g., Krause et al., 2019; Reato et al., 2013) and computational studies (Ali et al., 2013; Huang et al., 2021; Reato et al., 2010) to suggest that during stimulation (i.e., online), endogenous oscillatory activity may be entrained to match the frequency and phase of tACS. Furthermore, phase-locking of endogenous oscillations to tACS has also been demonstrated in human studies using electroencephalography (EEG; Helfrich et al., 2014a,b) and magnetoencephalography (MEG; Witkowski et al., 2016). Importantly, this online entrainment is thought to be mediated by resonance dynamics between the endogenous oscillations and exogenous current that are characterised by a phenomenon referred to as the “Arnold Tongue”, where oscillations that deviate from the eigenfrequency (i.e., resonant frequency) of the network require greater current intensities in order to be entrained by tACS (Ali et al., 2013; Huang et al., 2021; Liu et al., 2018; Schutter and Wischnewski, 2016; Thut et al., 2017; Vosskuhl et al., 2018).

Apart from the online effects observed during stimulation, many studies have also reported significant offline effects that are observed post-stimulation (for review see Veniero et al., 2015). Some studies have reported evidence suggesting that online entrainment by rhythmic stimulation may persist beyond the stimulation period (Hanslmayr et al., 2014; Marshall et al., 2006; Thut et al., 2011; van Bree et al., 2021), and as a result, these offline effects have traditionally been viewed as “echoes” of online entrainment. However, these entrainment echoes reportedly remain stable for a maximum of only a few oscillatory cycles (i.e., up to a few seconds) following cessation of stimulation and are therefore unlikely to be sufficient to explain offline tACS effects that are observed well beyond the proposed duration for entrainment echoes (herein referred to as aftereffects; Kasten et al., 2016; Neuling et al., 2013). Furthermore, aftereffects on oscillatory power have been reported even when stimulation was applied with a deliberate phase-shift to disrupt phase-continuity between successive tACS trials; (i.e., when optimal entrainment conditions were explicitly disrupted; Vossen et al., 2015). Therefore, the aftereffects of tACS have been primarily attributed to mechanisms related to neuroplasticity.

On the contrary, however, there is also some evidence to suggest that entrainment and neuroplastic aftereffects induced by tACS may not be mutually exclusive. For example, Helfrich et al. (2014a,b) reported a positive relationship between the magnitude of online entrainment and the magnitude of offline aftereffects. Although the magnitudes of these two effects were positively correlated, the authors further reported that entrainment was restricted to a narrow band around the peak frequency, whilst aftereffects were less frequency-specific and were spread across a wider band around the peak frequency, supporting the notion that the mechanisms underlying these effects are at least partially dissociable (Veniero et al., 2015; Vossen et al., 2015). Therefore, although online effects can likely be attributed to entrainment, offline aftereffects may instead reflect entrainment-mediated changes in synaptic plasticity.

Human studies investigating the mechanisms of tACS have traditionally used EEG (Helfrich et al., 2014a,b) or MEG (Witkowski et al., 2016) to quantify the online and offline effects of tACS on endogenous neural activity. However, the presence of complex, non-linear tACS artefacts in the EEG/MEG signal makes definitively demonstrating entrainment in the EEG/MEG concurrently with tACS a problematic task (Kasten and Herrmann, 2019; Noury et al., 2016; Noury and Siegel, 2017). Although many analysis methods have been developed in an attempt to filter the tACS artefact from the EEG/MEG, none of these approaches have been successful in eliminating the artefact in its entirety (for review see Kasten and Herrmann, 2019). Therefore, it is theoretically possible that previously reported entrainment effects in the EEG/MEG (Helfrich et al., 2014a; Witkowski et al., 2016) may reflect these residual artifacts rather than genuine entrainment of endogenous oscillations.

We have recently utilised a phase-dependent transcranial magnetic stimulation (TMS) protocol (Raco et al., 2016; Schaworonkow et al., 2018; Schaworonkow et al., 2019; Zrenner et al., 2018) as an alternative method for assessing entrainment to slow-wave tACS (Geffen et al., 2021). Here, TMS pulses are applied at different phases of tACS such that the TMS-induced motor evoked potential (MEP) amplitudes (assessed via electromyography; EMG) provide an indirect assessment of corticospinal excitability (CSE; Bergmann et al., 2012; Di Lazzaro et al., 2004; Hallett, 2007; Ilmoniemi and Kicić, 2010) across tACS phase. Phase-dependent TMS can therefore be used to assess whether CSE, and in turn, endogenous neural activity is entrained with respect to tACS phase. Crucially, unlike EEG, TMS–EMG measures are free of tACS artefacts, thus providing an unambiguous assessment of tACS-induced entrainment.

Although we did not observe entrainment of endogenous slow (0.75 Hz) oscillations in our previous experiment (Geffen et al., 2021), we attributed this null result to a lack of resonance dynamics between the eigenfrequency and stimulation frequency (Ali et al., 2013) as well as the relatively small MEP sample size per participant compared to simulation studies (Zoefel et al., 2019). The current study aimed to address both these limitations by using a tailored stimulation frequency that matches the eigenfrequency of the participants’ sensorimotor cortex during wake, and by performing multiple experiment sessions for each participant then pooling the data across sessions.

Mu (µ) oscillations (8–13 Hz) are one of the most prominent rhythms in the sensorimotor cortex during resting wake (Craddock et al., 2017; Hari, 2006; Weisz et al., 2014; Zhang and Ding, 2010) and are proposed to modulate CSE in a phase-specific manner (Baur et al., 2020; Berger et al., 2014; Bergmann et al., 2019; Hussain et al., 2018; Schaworonkow et al., 2018; Schaworonkow et al., 2019; Stefanou et al., 2018; Wischnewski et al., 2022; Zrenner et al., 2018;), with µ troughs thought to reflect active facilitation of gamma activity and information processing, referred to as the “pulsed facilitation” hypothesis (Bergmann et al., 2019). However, these troughs may instead reflect “pulsed inhibition” (akin to occipital alpha oscillations; Mathewson et al., 2011; Mazaheri and Jensen, 2010; Schalk, 2015) of the primary somatosensory cortex (S1) that results in a net transient local disinhibition of feed-forward inhibitory inputs from S1 to the primary motor cortex (M1; Bergmann et al., 2019; Murray and Keller, 2011; Thies et al., 2018; Turco et al., 2018). Therefore, it is still not clear whether µ activity in the sensorimotor cortex reflects active facilitation, inhibition, or possibly even a symmetric combination of the two (Bergmann et al., 2019).

The phase-specificity of the µ rhythm has been further supported by evidence that TMS-induced long-term potentiation-like plasticity occurs preferentially when TMS pulses are applied during µ troughs whilst long-term depression-like plasticity occurs during peaks (Baur et al., 2020; Zrenner et al., 2018). Furthermore, Hussain et al. (2021) demonstrated greater broadband oscillatory power responses and improved offline motor learning following TMS at µ troughs but not at µ peaks.

Whilst the number of studies assessing the effects of µ-tACS on CSE are limited, recent studies report a facilitation of MEP amplitudes from µ-tACS (Feurra et al., 2019; Fresnoza et al., 2018; Madsen et al., 2019), supporting a facilitatory role of µ oscillations on CSE (Bergmann et al., 2019; Karabanov et al., 2021; Ogata et al., 2019; Thies et al., 2018; Wischnewski et al., 2022), whether that be via active facilitation of M1 or active inhibition of S1 to disinhibit M1. However only one of these studies (Madsen et al., 2019) assessed the relationship between µ-tACS phase and CSE, finding no effect of phase on CSE. Therefore, there remains a crucial need for further investigation of both the acute phasic effects and sustained aftereffects of µ-tACS on CSE.

### Aims

The aims of this experiment were (1) to investigate phasic entrainment of CSE to the phase of intermittent µ-tACS, both during (i.e., online entrainment) and immediately following (i.e., entrainment echoes) stimulation; (2) to investigate the sustained aftereffects of µ-tACS on CSE and µ EEG power.

### Hypotheses

It was hypothesised that µ-tACS would induce µ rhythm-like sinusoidal changes in CSE that correspond with the tACS phase, demonstrating online entrainment. Second, that sinusoidal changes in CSE would persist for the first oscillatory cycle immediately following each trial of stimulation, thus demonstrating entrainment echoes. Third, that CSE and µ EEG power would increase across the total stimulation period, and these increases would then be sustained beyond the stimulation period (i.e., plastic aftereffects).

## 2. Materials and Methods

### 2.1. Subjects

Thirteen neurotypical, right-handed participants were recruited for the experiment (5 male, mean age = 23 ± 4 years). Each participant completed three separate experimental sessions (with a minimum of 1 week between sessions) to maximise the MEP sample size per participant for assessing phasic entrainment of MEP amplitudes. The MEP sample size (rather than the participant sample size) is the most important factor for increasing the sensitivity of phasic analyses (Zoefel et al., 2019). All participants completed a safety screening questionnaire (Keel et al., 2001) and provided a written statement of informed consent prior to commencing the experiment. The exclusion criteria from this safety screen were identical to our most recent experiment (Geffen et al., 2021) and included: “personal or family history of epilepsy/seizures, medication that could affect seizure threshold, history of brain injury/condition (e.g., stroke, concussion, etc.), implanted devices or metal in the head, frequent or severe headaches, or current pregnancy”. For the duration of the experiment sessions, participants were instructed to relax, avoid moving their hands, and avoid doing any mentally stimulating tasks in their head (e.g., simple math) but also to keep their eyes open and stay awake. Approval was granted by The University of Queensland Human Research Ethics Committee. Experimental sessions took place in the Brain Stimulation Laboratory of the School of Health and Rehabilitation Sciences at The University of Queensland.

### 2.2. Experimental Setup

#### 2.2.1 tACS/TMS

The target site for stimulation was the hand area of the left primary motor cortex (M1_HAND_; typically corresponds with the EEG coordinate C3), specifically the region associated with the right first dorsal interosseous (FDI; primary target muscle) and abductor digiti minimi (ADM; secondary target muscle), intrinsic muscles in the index- and little-finger respectively.

TMS pulses were applied via a Magstim Double 70mm Remote Control Coil charged by a Magstim 200^2^ stimulator (Magstim, UK). Manual TMS “hot-spotting” (Rossini et al., 1994) was performed to determine each participant’s individual location for the left M1_HAND_ region as well as the TMS intensity required to consistently induce MEPs in the FDI (i.e., our primary target muscle) with amplitudes of ∼1 mV (Cuypers et al., 2014; Ogata et al., 2019; Thies et al., 2018). Here, the position of the TMS coil and the stimulation intensity are systematically adjusted until MEPs are consistently induced with amplitudes around the target amplitude. For some participants (3 total, not included in the reported sample size), MEPs of this amplitude could not be consistently induced during hot-spotting at intensities <80% of the maximum stimulator output, and so these participants did not take any further part in the study. This threshold was set to avoid overheating of the TMS coil mid-session at intensities above 80% of maximum stimulator output. The location of the left M1_HAND_ region was marked on the participant’s scalp using an erasable marker.

tACS was applied via a NeuroConn DC Stimulator Plus using a classical M1-contralateral supraorbital region electrode montage (Heise et al., 2016), with the target electrode placed ∼2 centimetres posterolateral to the marked hotspot (around Cp3) and the return electrode placed over the contralateral supraorbital region. This slight posterolateral shift for the M1 electrode is thought to enhance the effectiveness of tACS by reducing current shunting through the scalp and cerebrospinal fluid to maximise the current density at M1 (Faria et al., 2011). The scalp was first rubbed with ethanol (70%), then Ten20 conductive paste (Weaver and Company) was applied to both 42×45mm pad electrodes before placing them onto their respective positions on the scalp. The cable for the M1 electrode was oriented inferiorly, whilst the cable for the supraorbital region electrode was oriented laterally.

#### 2.2.2. EMG/EEG

EMG activity for the FDI and ADM muscles was recorded via disposable surface electrodes (H124SG 30mm x 24mm). For both muscles, one electrode was placed on the relevant muscle belly and the other on the nearby metacarpophalangeal joint, with reference electrodes placed over the palmar aspect of the distal forearm.

EEG activity was recorded using 66 reusable surface electrodes (64 active electrodes in 10:20 layout + 2 reference electrodes) embedded in a cap. The EEG electrodes directly above the tACS pad electrodes had to be removed as the current output at these pads exceeds the acceptable limit of the EEG system, preventing EEG data acquisition. The specific EEG channels that needed to be removed varied slightly between participants due to interindividual differences in tACS electrode placement. For the tACS pad targeting M1_HAND_, EEG channels C1 and C3 were removed for all sessions, channels Cp1 and Cp3 were removed for all but one participant, and channels Fc1 and Fc3 were removed for 8 of the sessions (across 4 participants). For the tACS pad over the contralateral supraorbital region, removed channels included Fpz, Fp2, Af4, and/or Af8.

Additional TMS hot-spotting was performed following the addition of the EEG cap, as the cap obscures the original marked hotspot whilst also slightly increasing the distance between the scalp and the TMS coil, and in most cases, the TMS intensity required to consistently induce MEPs around the target amplitude (see Farzan et al., 2016). This increased distance between the scalp and coil was minimised by removing the plastic wells for the EEG electrodes directly above the M1 tACS pad electrode. The revised hotspot was marked on the EEG cap using tape and an erasable marker.

#### 2.2.3. Data Collection

EEG electrode measurements were captured (2048 Hz sampling rate) using a Biosemi ActiveTwo AD-Box (Biosemi, Amsterdam, Netherlands). EMG electrode measurements for both the FDI and ADM were acquired (1 kHz sampling rate, 20–1000 Hz band pass filtering) via an electrode adaptor (Model CED1902), before being amplified by a CED1902, and finally recorded by a CED1401 MICRO3 (Cambridge Electronic Designs, Cambridge, UK). TMS triggers were directly recorded by the CED1401 MICRO3 and were also recorded by the ActiveTwo EEG system via an MMBT-S Trigger Interface Box. The EMG data were then transferred from the CED1401 MICRO3 to a PC and saved via Signal software (6.04; Cambridge Electronic Designs, Cambridge, UK), whilst the EEG data were transferred from the ActiveTwo AD-Box to a separate PC and saved via ActiView (Ver. 8.08) software (Biosemi, Amsterdam, Netherlands). Both EMG and EEG data were then exported to MATLAB (Ver. R2020b) and subsequently JASP (Ver. 0.14.1.0) for analysis.

### 2.3. Experimental Procedure

#### 2.3.1. EEG Paradigm

Before commencing the experiment, EEG activity was recorded for 5-minutes of wakeful (eyes open) rest and analysed using a Welch’s power spectral density estimate to determine each participant’s individual µ frequency (IMF), which was used as the stimulation frequency for tACS. The IMF was defined objectively as the frequency with the greatest spectral power from 8–13 Hz (at a resolution of 0.5 Hz), even if the frequency spectrum included multiple peak frequencies with similar power values (e.g., from a broad peak around a particular frequency or from multiple peaks across different frequencies). Additional 5-minute recordings were made during two breaks within the stimulation period as well as at the end of the experimental session to assess any changes in µ (8–13 Hz) and IMF (± 1 Hz) power. EEG was also recorded throughout the tACS delivery to provide a measure of the tACS output, which allowed us to determine when TMS pulses were applied relative to the tACS phase, as well as to record resting EEG data for the echo periods.

#### 2.3.2. tACS Paradigm

To desensitise the participants to the tACS sensations, each participant received an initial “test” tACS paradigm after the electrodes were attached, consisting of 30 seconds of tACS at an intensity of 2 mA and a generic frequency of 10 Hz with an additional 5 seconds of ramp up/down at the start and end of the stimulation paradigm. For the actual tACS paradigm (Figure 1), participants received 90 trials of tACS in each experiment session, with each trial consisting of 16 seconds of tACS (“online”) at an intensity of 2 mA and a frequency that was tailored to the participant’s IMF, followed by 8 seconds of no tACS (“offline”), for a total of 24 minutes of tACS and 12 minutes of no tACS (Fig 1A). The entire stimulation period was divided into 3 blocks (12 minutes comprising 30 trials each), with 5-minute break periods (no stimulation delivered) between blocks. This allowed us to record continuous periods of EEG activity within the stimulation session without the presence of tACS artifacts, whilst also allowing the TMS coil to cool.

**Figure 1.**
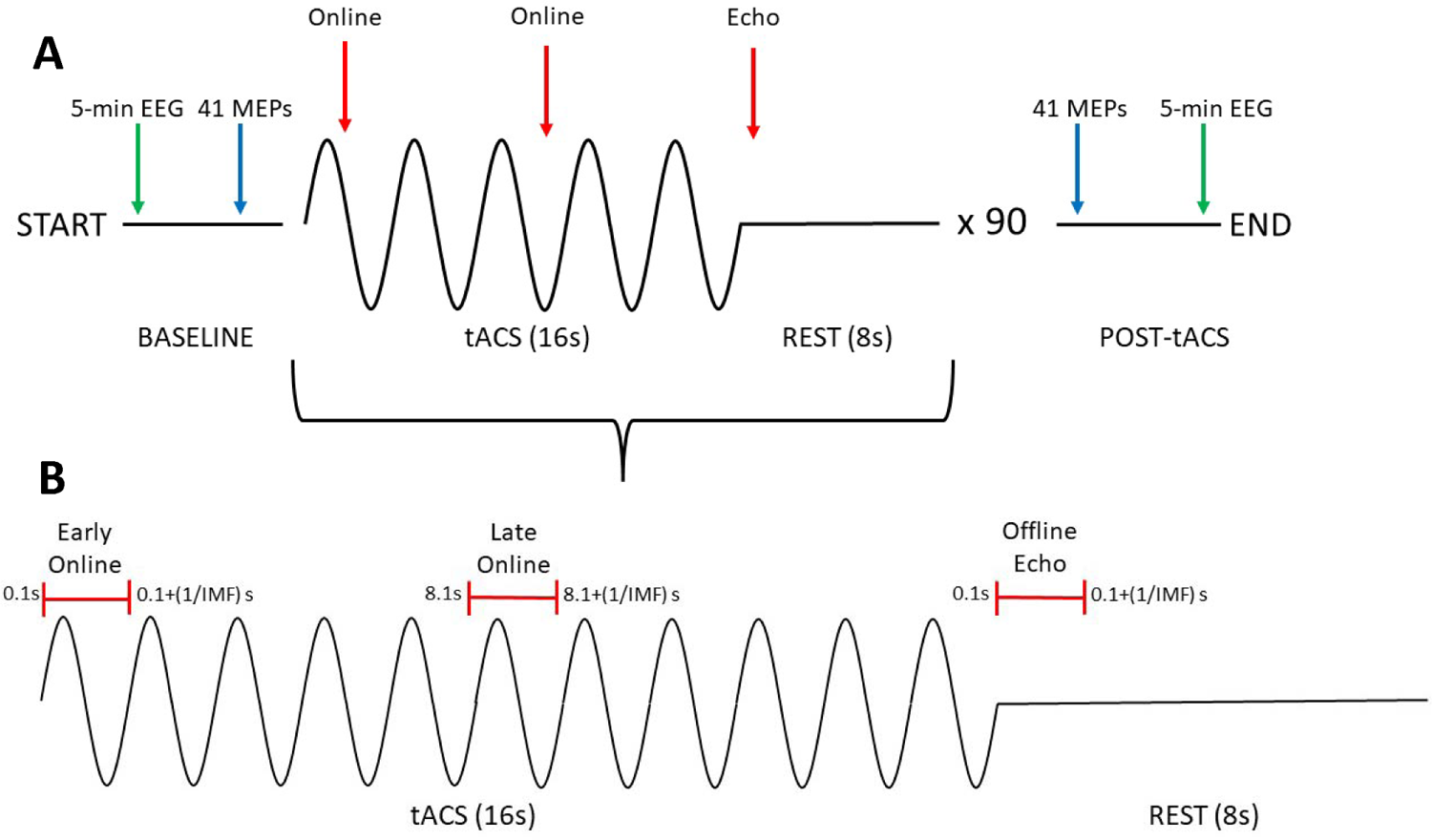
Experimental procedure for probing changes in CSE induced by µ-tACS. A) The experimental session consisted of a 16-second tACS period (represented as sine waves) followed by an 8-second rest period (represented as flat lines), repeated 90 times for a total of 36 minutes. Resting EEG activity was recorded for 5-minutes (green arrows) at the start of the session to determine each participant’s IMF, as well as at two 5-minute breaks (at 12- and 24-minutes, not shown in figure) and at the end of the session to assess any tACS-induced changes in oscillatory activity. To assess the aftereffects of µ-tACS on CSE, 41 TMS-induced MEPs were acquired at the start and the end of the experimental session (blue arrows). To assess the phasic effects of µ-tACS on CSE online to and immediately following tACS, 2 MEPs were acquired within each tACS period and 1 MEP was acquired within each rest period (red arrows). B) To probe the phasic effects of µ-tACS on CSE (both online and offline), MEPs were acquired at 3 epochs within each tACS trial (1 MEP per epoch per trial, 90 MEPs per epoch per session, 270 MEPs per epoch across the 3 sessions): early online, late online, and offline echo. TMS pulses were applied with a “jitter” (i.e., randomised time delay equal) that covers the length of a single tACS cycle (i.e., 1/IMF) so that pulses are approximately uniformly delivered across the different phases of tACS.

#### 2.3.3. TMS Paradigm

TMS pulses were applied at 3 epochs within each trial (1 MEP per epoch per trial, 90 MEPs per epoch per session, 270 MEPs per epoch across the 3 sessions): early online (∼0.1–0.2s after tACS starts), late online (∼8.1–8.2s after tACS starts), and offline echo (∼0.1–0.2s after tACS ends).

To examine the phasic effects of µ-tACS on CSE (both online and offline), sufficient MEPs need to be acquired across the different phases of the tACS. This was achieved by implementing a “jitter” (i.e., a randomised time delay that covers the length of a single tACS cycle; see Fig 1B) to the delivery of TMS so that it is not locked to a specific phase of tACS, and thus, TMS pulses are approximately uniformly delivered across the phase of the tACS cycle.

To examine the aftereffects of the entire tACS paradigm on CSE, 41 MEPs were also acquired at baseline and at the end of the entire period of tACS delivery (delivered at ∼0.2 Hz). For sessions where the TMS intensity approached the upper limit for acceptable coil temperatures (∼80% intensity), the coil needed to be cooled for 2 minutes in between the tACS offset and post-tACS MEP acquisition.

### 2.4. Statistical Analysis

#### 2.4.1 Data Transformation

The EEG data were imported to MATLAB (Ver. 2020b) via the EEGLAB toolbox (Delorme and Makeig, 2004). The data were bandpass filtered (1–40 Hz) then artefacts (e.g., eye blinks) and bad channels (i.e., channels with a flat or noisy signal) were automatically removed. Finally, channels Fc1, Fc3, Fc5, C5, and Cp5 (as well as Cp1 and Cp3 for the one participant that did not have these channels removed) were analysed using a Welch’s power spectral density estimate and subsequently log transformed to determine each participant’s IMF as well as their spectral power from 8–13 Hz and from (IMF-1)–(IMF+1) Hz. Relative µ/IMF spectral powers were then computed by subtracting the lowest power value from all values (to remove any negative values caused by the log transform) then dividing all power values by the sum of powers for all frequencies.

Due to an indexing error in the MATLAB script for the spectral analysis that was not discovered until after data collection was completed, the EEG channels that were analysed for the initial IMF estimates varied slightly from the desired channels (i.e., the ones listed above) for some of the sessions. This indexing error was related to the “clean_rawdata” function (EEGLAB toolbox) that automatically removes artifacts and bad channels from the data, which had caused the remaining clean channels to be assigned to different channel numbers, thus changing the channels that were selected for spectral analysis. It should also be noted that for the initial IMF estimates, the EEG data were re-referenced to the common average; however, subsequent analyses were performed on data using the “native” EEG reference electrodes, since the common average referencing resulted in subtle artefacts that interfered with the echo and break period spectral analyses. After interpolating the removed channels and reperforming the spectral analyses using the correct channels (i.e., Fc1, Fc3, Fc5, C5 and Cp5, as well as Cp1 and Cp3 for the one participant that did not have these channels removed) and the native reference electrodes, it was found that the revised IMF estimates (i.e., the endogenous eigenfrequency) differed from the initial IMF estimates (i.e., the stimulation frequency) for 17 out of 39 sessions (8 from the indexing error and 9 from the change in referencing), with the differences between the initial and revised estimates ranging from 0.5–3 Hz. These changes to the indexing and referencing were subsequently applied for the remaining EEG analyses, although it should be noted that the spectral power analyses for IMF ± 1 Hz were performed with respect to the initial IMF estimates rather than the revised IMF estimates, since tACS-induced entrainment is expected to occur specifically with respect to the stimulation frequency rather than the endogenous eigenfrequency (Ali et al., 2013). Although we initially considered discarding the incorrectly targeted sessions, we felt that the inclusion of these data could provide valuable information regarding the necessity to accurately target the IMF. Therefore, we have included all sessions in the analysis (described below).

The MEP exclusion criteria were identical to those used in our previous experiment (Geffen et al., 2021) and included: “the first MEP for each data set (as well as the first MEP after each of the break periods) which may be larger (Brasil-Neto et al., 1994) and more variable (Schmidt et al., 2009) than subsequent MEPs, as well as any MEPs with voluntary EMG activity detected in the 500 ms prior to TMS delivery (∼1–2% of MEPs excluded).”

The TMS triggers were automatically categorized into their respective epochs (i.e., early online, late online, and offline echo) and the tACS phase was calculated using the EEG data from channel Oz around the “late online” triggers, since these triggers are the only ones where tACS was present both before and after TMS was applied, thus, providing the most reliable estimate of tACS phase. The computed phase for the late online triggers was then extrapolated (both forward and backward) and its values computed at each of the other epochs, since we expect the tACS phase to continue into the offline period if entrainment persists beyond stimulation.

#### 2.4.2. Data Analysis

To assess phasic entrainment of MEP amplitudes with respect to tACS phase, a permutation analysis (Zoefel et al., 2019) was performed. Here, an ideal (best-fitting) sinusoidal model is fitted to each participant’s observed MEP amplitudes (∼270 MEPs pooled across 3 sessions for each epoch) based on their tACS phase (Bland & Sale, 2019). The MEP amplitudes are then shuffled with respect to their phases for a total of 1000 permutations per participant and ideal sinusoidal models are fitted to the shuffled data. The true and shuffled sinusoidal model amplitudes are then compared, with the individual P-values representing the proportion of shuffled model amplitudes exceeding the true model amplitudes, remembering that under the null hypothesis (which assumes that the observed effects are not phase-specific) the amplitudes of the sinusoidal models should be small (i.e., closer to zero). Because the permutation procedure disrupts any phasic effects that may be present, the shuffled MEPs act as a negative control for the true MEPs, and thus, the permutation analysis does not require a sham stimulation condition as a negative control. The group P-value for each muscle/timepoint was then obtained by combining the individual P-values using Fisher’s method (Zoefel et al., 2019).

To determine if there was a significant difference in mean MEP amplitudes between the pre- and post-tACS MEPs (41 each per session, 123 across the 3 sessions), two-way repeated measures ANOVAs (rmANOVA) were performed with STIMULATION (pre, post) and SESSION (1, 2, 3) as the two repeated measures factors (to confirm that there were no significant differences between the three sessions). To determine if there was a significant difference in mean MEP amplitudes between the online MEPs (540 across the 3 sessions) and the offline echo MEPs (270 across the 3 sessions) or between the three tACS blocks (180 online MEPs and 90 offline echo MEPs per block across the 3 sessions), a three-way repeated measures ANOVA was performed with STIMULATION (online, offline), BLOCK (1, 2, 3), and SESSION (1, 2, 3) as the three repeated measures factors. Post hoc t-tests (corrected for multiple comparisons using Holm’s method) were then performed to compare the individual groups against each other. To examine how offline changes in MEP amplitudes evolve throughout the tACS period, mean MEP amplitudes for the pre- and post-tACS MEPs (123 each across the 3 sessions) and the offline MEPs of each tACS block (90 per block across the 3 sessions) were compared using a two-way repeated measures ANOVA with TIME (5 levels: pre-tACS, post-tACS, and the offline MEPs for the 3 tACS blocks) and SESSION (1, 2, 3) as the two repeated measures factors. Again, post hoc t-tests were then performed to compare the individual groups against each other. To examine the effects of the tACS paradigm on both µ (8-13 Hz) and IMF (± 1 Hz) relative spectral power, a two-way repeated measures ANOVA was performed with TIME (7 levels: pre-tACS, post-tACS, the 2 break periods, and the pooled echo periods for each of the 3 tACS blocks) and SESSION (1, 2, 3) as the two repeated measures factors. Again, post hoc t-tests were then performed to compare the individual groups against each other. For the repeated measures ANOVAs, standardized effect sizes for any significant differences were calculated as η^2^ values. For the post hoc t-tests, standardized effect sizes for any significant differences were calculated as Cohen’s d values.

We later performed separate analyses for the sessions that showed a difference between initial and revised IMF estimates (henceforth referred to as “affected” sessions) and the sessions that did not show a difference (henceforth referred to as “unaffected” sessions). However, because the number of affected and unaffected sessions varied between participants, most of these separate analyses had to be performed using data that had been averaged across the affected/unaffected sessions for each participant rather than including SESSION as a repeated measures factor (except for the permutation analysis where the data was simply pooled across the affected/unaffected sessions rather than being pooled across all sessions).

## 3. Results

### 3.1. Phase-Specific Effects of µ-tACS on MEP Amplitude

The acute phase-specific effects of µ-tACS on MEP amplitudes were assessed by a permutation analysis. Ideal sinusoidal models were fitted to each participant’s observed MEP amplitudes based on their tACS phase for each epoch (∼90 MEPs per epoch per session, ∼270 MEPs per epoch per participant, see Figure 2) and the amplitudes of these models were compared to the model amplitudes of 1000 permutations of the MEPs. The individual and group P-values for each muscle and epoch are listed in Table 1 and summarised in Supplementary Figure 1. Fitted sinusoidal models with P < 0.05 are shown in Figure 2.

**Figure 2.**
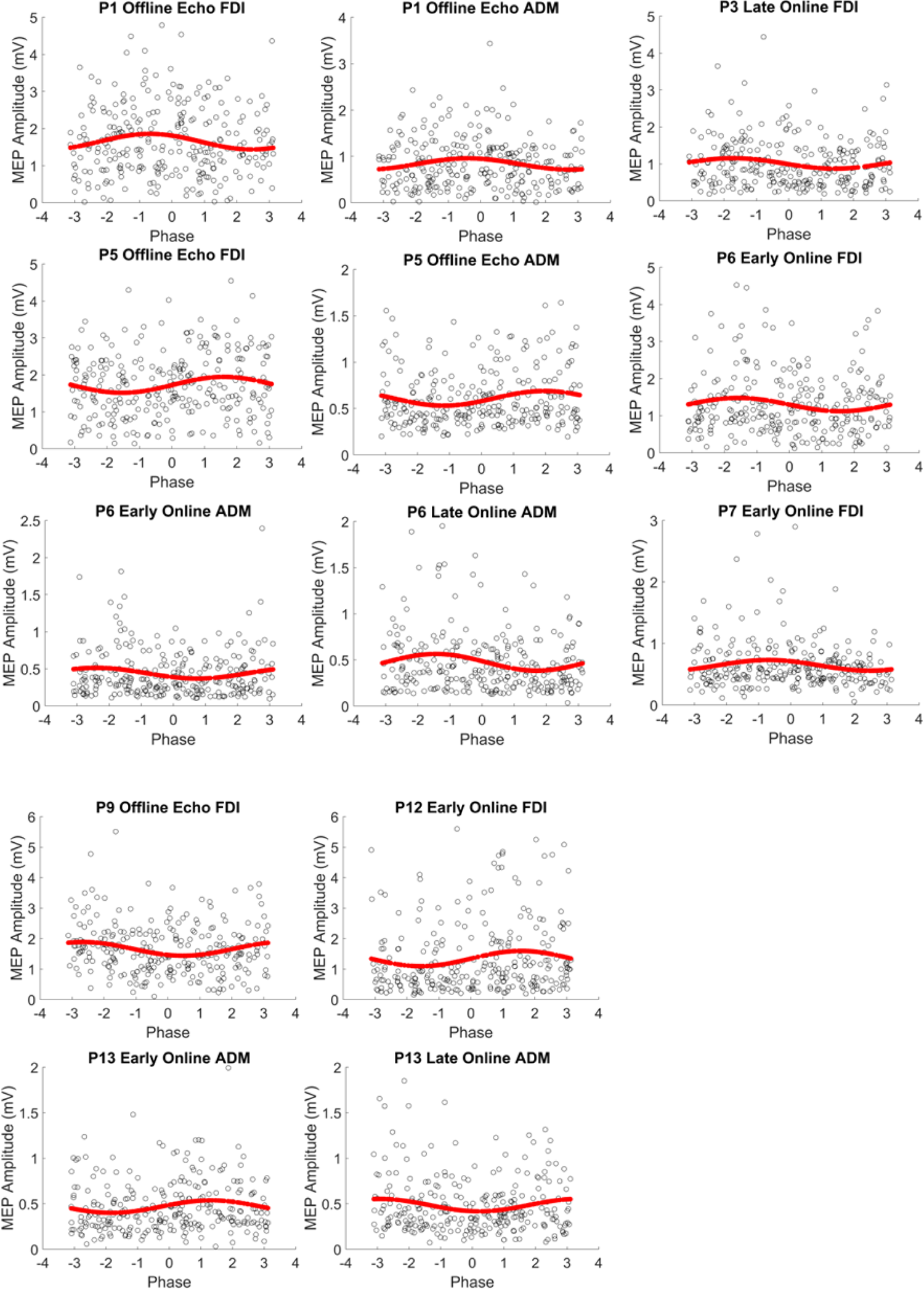
Participant’s Fitted Sinusoidal Models with Significant Entrainment to tACS phase. For each plot, markers represent the observed MEP amplitudes for the participant (e.g., P1 denotes Participant 1), epoch (Early Online, Late Online, or Offline Echo), and muscle (FDI or ADM) denoted above the plot (∼270 MEPs across 3 sessions) sorted according to tACS phase. A phase value of 0 denotes the tACS peak, π or -π denotes the trough, π/2 denotes the falling edge, and – π/2 denotes the rising edge. Red lines represent the fitted sinusoidal models for these MEPs, which had greater amplitudes than the fitted sinusoidal models for at least 950 out of 1000 permutations of the MEPs (i.e., P < 0.05).

**Table 1.**
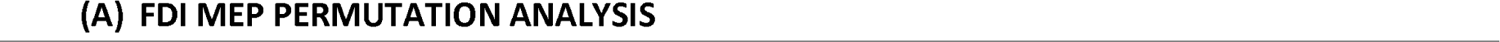
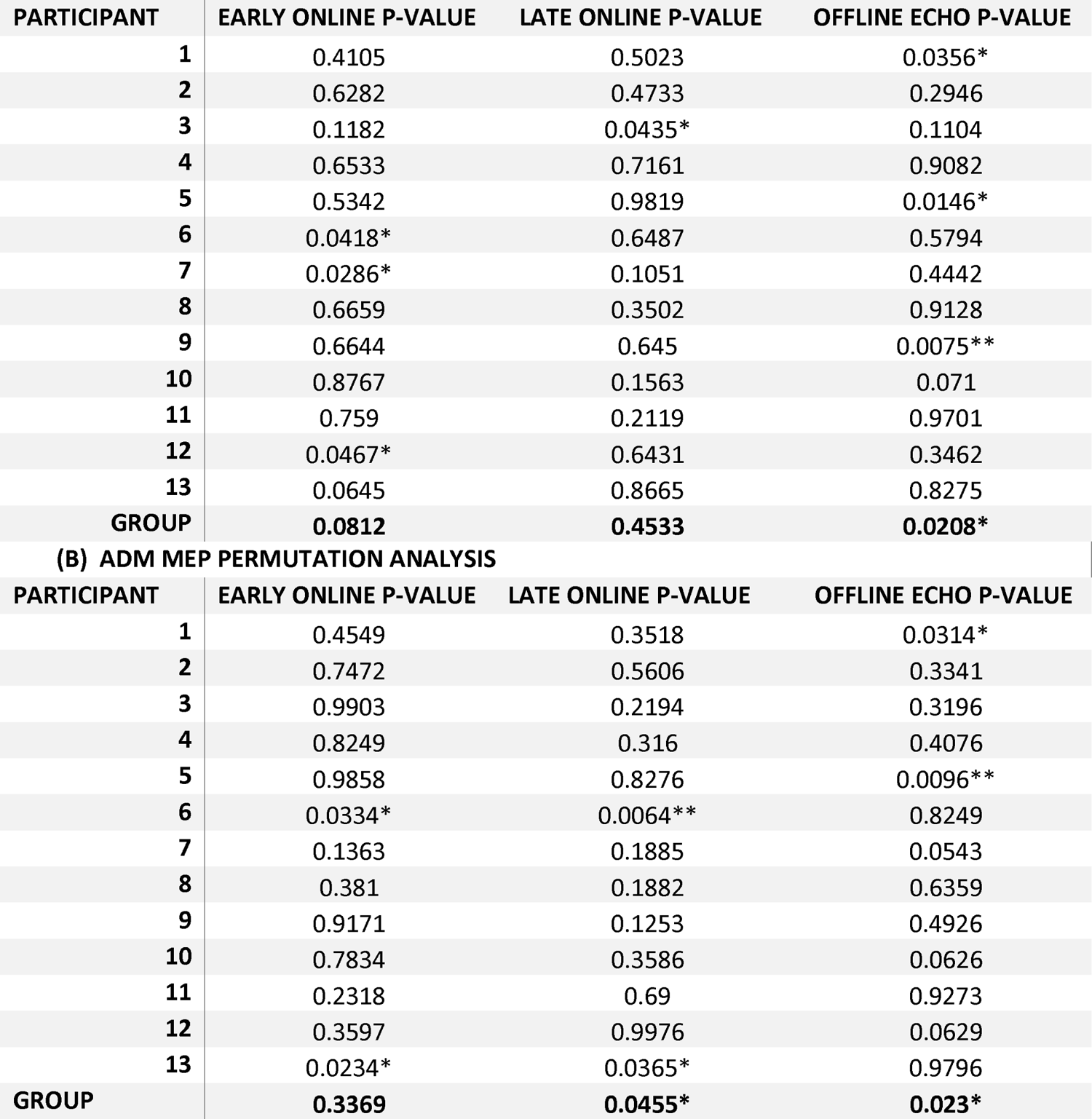
Table of Individual and Group P-Values for Permutation Analysis of Phasic tACS Effects on FDI (A) and ADM (B) MEP Amplitude. P-values > 0.05 are denoted by * whilst P-values > 0.01 are denoted by **. (A) Participants 6, 7, and 12 showed significant phasic entrainment of FDI MEP amplitudes for the early online MEPs. Participant 3 showed significant entrainment for the late online MEPs. Participants 1, 5, and 9 showed significant entrainment for the offline echo MEPs. Revealed significant entrainment for the offline echo MEPs (P = 0.0208) but no entrainment for either of the online MEPs, although the early online MEPs approached significance (P = 0.0812). (B) Participants 6 and 13 showed significant phasic entrainment of ADM MEP amplitudes for both early and late online MEPs; Participants 1 and 5 showed significant entrainment for the offline echo MEPs. The group analysis revealed significant entrainment for the offline echo MEPs (P = 0.023) as well as the late online MEPs (P = 0.0455) but no entrainment for the early online MEPs. N = 13

#### 3.1.1. Target muscle (FDI) MEP amplitudes

As shown in Table 1A, participants 6, 7, and 12 showed significant phasic entrainment of FDI MEP amplitudes for the early online MEPs. Participant 3 showed significant entrainment for the late online MEPs. Finally, participants 1, 5, and 9 showed significant entrainment for the offline echo MEPs. The group analysis revealed significant entrainment for the offline echo MEPs (P = 0.0208) but no entrainment for either of the online MEPs, although the early online MEPs approached significance (P = 0.0812).

#### 3.1.2. Non-target muscle (ADM) MEP amplitudes

As shown in Table 1B, participants 6 and 13 showed significant phasic entrainment of ADM MEP amplitudes for both early and late online MEPs whilst participants 1 and 5 showed significant entrainment for the offline echo MEPs. Similar to the FDI, the group analysis for the ADM revealed significant entrainment for the offline echo MEPs (P = 0.023). Furthermore, there was also significant entrainment for the late (but not early) online MEPs (P = 0.0455).

### 3.2. Aftereffects of µ-tACS on MEP Amplitude

The aftereffects of the tACS paradigm on CSE were assessed by comparing mean MEP amplitudes pre- and post-tACS using a two-way rmANOVA. As shown in Figure 3, MEP amplitudes were found to be significantly greater post-tACS compared to pre-tACS for both the FDI (Left; mean_pre_ = 0.97 mV ± 0.17; mean_post_ = 1.39 mV ± 0.5; F_1,12_ = 16.02, p = 0.002, η^2^ = 0.167; **) and ADM (Right; mean_pre_ = 0.46 mV ± 0.17; mean_post_ = 0.69 mV ± 0.29; F_1,12_ = 27.11, p < 0.001, η^2^ = 0.257; ***). Furthermore, there was no main effect of SESSION for either muscle (FDI: F_2,24_ = 0.884, p = 0.426; ADM: F_2,24_ = 0.805, p = 0.459) nor were there any STIMULATION × SESSION interactions (FDI: F_2,24_ = 0.181, p = 0.836; ADM: F_2,24_ = 0.404, p = 0.672), confirming that there were no significant differences in MEP amplitude between the three sessions.

**Figure 3.**
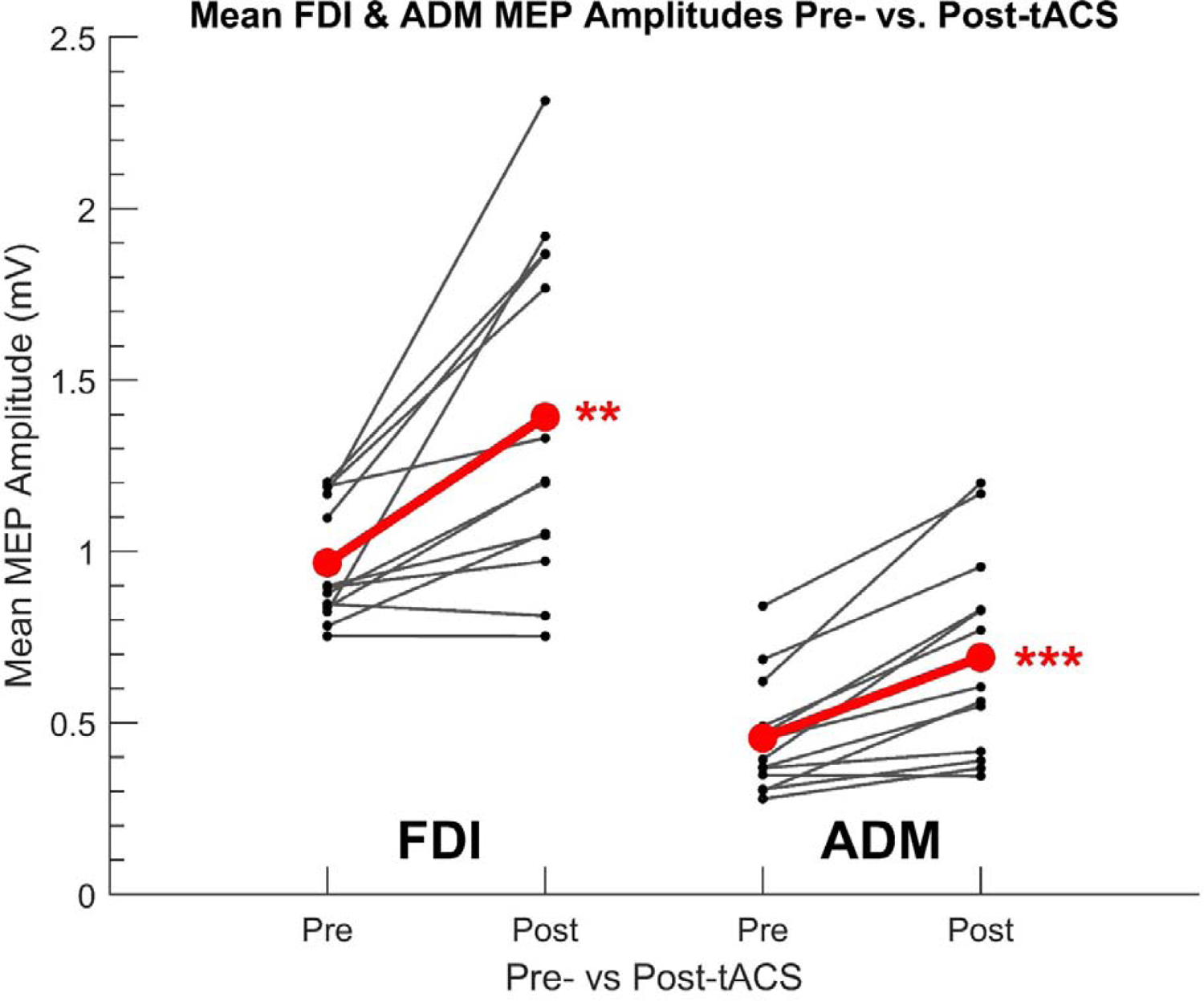
Individual Mean FDI (Left) and ADM (Right) MEP Amplitudes Before and After µ-tACS. Points represent each participant’s mean MEP amplitude from 120 TMS-induced MEPs per participant (40 MEPs per experiment session) acquired before (pre-tACS) and after (post-tACS) receiving 90 “trials” (∼36 minutes) of µ-tACS, with each trial consisting of 16 seconds of tACS followed by 8 seconds of rest. Individual differences in mean MEP amplitude are represented by the solid lines. The global mean MEP amplitude for each group is represented by a red point, with the global mean difference from pre- to post-tACS represented by a red line connecting the two red points. A significant increase in MEP amplitude from pre-tACS to post-tACS was reported for both the FDI (two-way rmANOVA: F_1,12_ = 16.02, p_12_ = 0.002, η^2^ = 0.167; **) and ADM (F_1,12_ = 27.11, p < 0.001, η^2^ = 0.257; ***). N = 13.

Mean MEP amplitudes from the offline periods of each block were then compared against each other as well as against the pre- and post-tACS mean amplitudes using a two-way rmANOVA. As shown in Figure 4, there was a significant main effect of TIME for both muscles (FDI: F_4,48_ = 8.125, p <0.001, η^2^ = 0.115; ADM: F_4,48_ = 10.828, p < 0.001, η^2^= 0.144); however, subsequent post-hoc t-tests revealed slight differences between the two muscles. The FDI (Figure 4A) showed significant increases in MEP amplitude between the pre-tACS MEPs and the offline MEPs from all 3 tACS blocks as well as the post-tACS MEPs (t_12_ = − 3.624, −3.469, −3.753, −4.002; p = 0.028, 0.032, 0.025, 0.018; d = −1.005, −0.962, −1.041, −1.110 for blocks 1, 2, 3, and post-tACS respectively) but no significant differences in MEP amplitude between the 3 blocks or between any of the tACS blocks and the post-tACS MEPs. This suggests a relatively sharp increase in FDI MEP amplitude within the 1^st^ tACS block that is then sustained throughout and beyond the stimulation period. Meanwhile, the ADM (Figure 4B) showed a significant increase in MEP amplitude between the pre-tACS MEPs and the MEPs from blocks 2 and 3 as well as the post-tACS MEPs (t_12_ = −4.231, −5.287, −5.207; p = 0.009, 0.002, 0.002, d = −1.173, −1.466, − 1.444 for blocks 2, 3, and post-tACS respectively) but no significant difference between pre-tACS and block 1 (t_12_ = −1.956; p = 0.258). There was, however, a significant increase from block 1 to block 2 (t_12_ = −4.255; p = 0.009; d = − 1.18) but no significant differences between blocks 1 and 3 (t_12_ = −3.112; p = 0.054) or blocks 2 and 3 (t_12_ = −2.084; p = 0.258). This suggests a more gradual increase in ADM MEP amplitude compared to FDI MEP amplitude. There were no main effects of SESSION for either muscle (FDI: F_2,24_ = 1.558, p = 0.231; ADM: F_2,24_ = 1.246, p = 0.306) nor were there any TIME × SESSION interactions (FDI: F_8,96_ = 1.247, p = 0.28; ADM: F_8,96_ = 1.922, p = 0.065), confirming that there were no significant differences in MEP amplitude between the three sessions.

**Figure 4.**
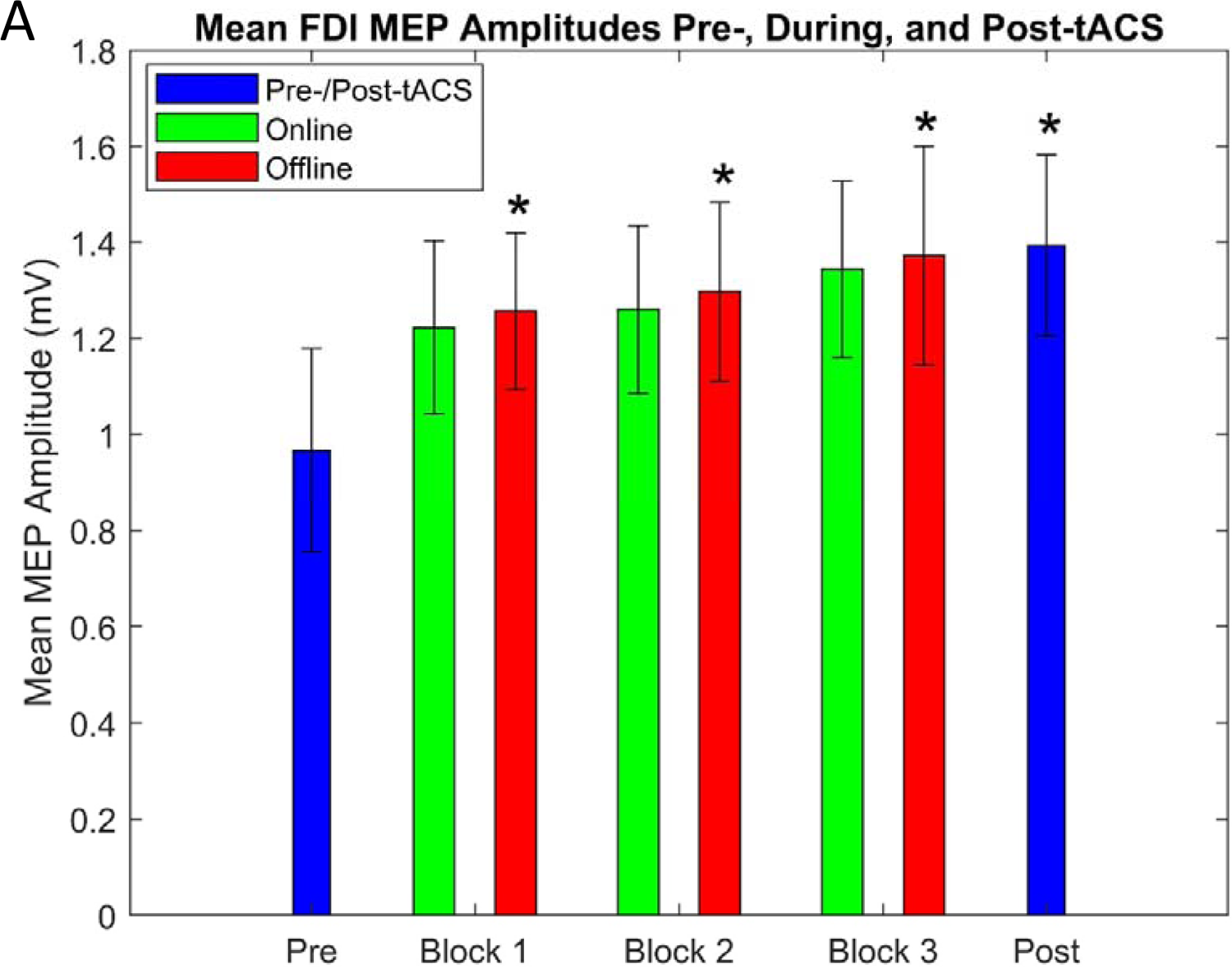
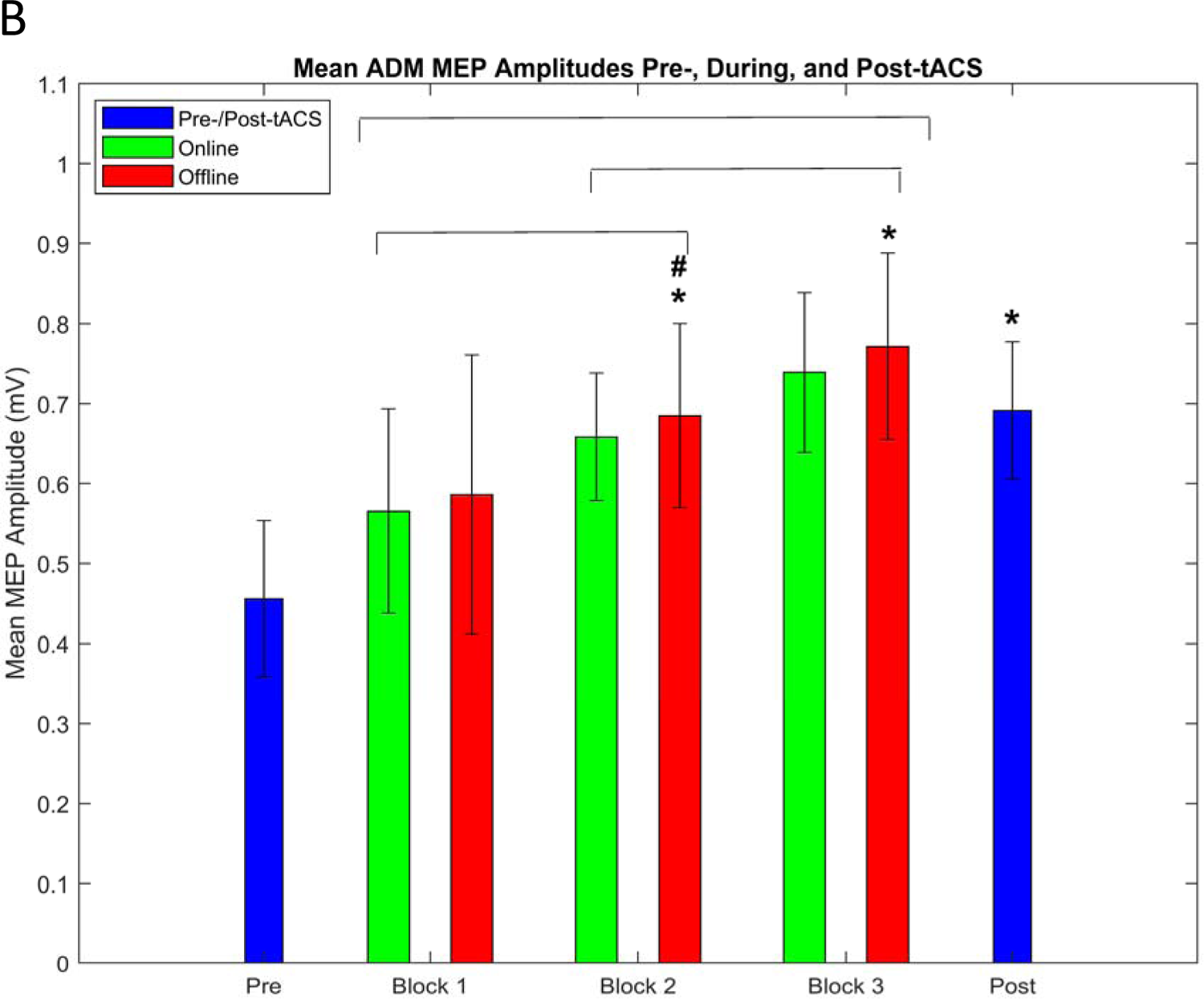
Group Mean FDI (A) and ADM (B) MEP Amplitudes Before, During, and After Receiving µ-tACS. Blue bars represent mean MEP amplitudes (with within-subjects error bars) from 40 TMS-induced MEPs (per participant) acquired before (pre-tACS) and after (post-tACS) receiving 90 “trials” (∼36 minutes, split into 3 30-trial “blocks”) of µ-tACS, with each trial consisting of 16 seconds of tACS (online) followed by 8 seconds of rest (offline). Mean MEP amplitudes from 60 MEPs acquired during the online periods and 30 MEPs acquired during the offline periods of each block are represented by green and red bars respectively. (A) There was a significant difference in FDI MEP amplitudes between the pre-tACS MEPs and the offline MEPs from all 3 tACS blocks as well as the post-tACS MEPs (t_12_ = −3.624, −3.469, −3.753, −4.002; p = 0.028, 0.032, 0.025, 0.018; d = −1.005, −0.962, −1.041, −1.110 for blocks 1, 2, 3, and post-tACS respectively; *). (B) There was a significant difference in ADM MEP amplitudes between the pre-tACS MEPs and the offline MEPs from blocks 2 and 3 as well as the post-tACS MEPs (t_12_ = −4.231, −5.287, −5.207; p = 0.009, 0.002, 0.002, d = − 1.173, −1.466, − 1.444 for blocks 2, 3, and post-tACS respectively; *) but no significant difference between pre-tACS and block 1 (t_12_ = −1.956; p = 0.258). There was also a significant difference in offline MEP amplitude between block 1 and 2 (t_12_ = −4.255; p = 0.009; d = −1.18; #) but no significant differences between blocks 1 and 3 (t_12_ = −3.112; p = 0.054) or blocks 2 and 3 (t_12_ = − 2.084; p = 0.258). When assessing both the online and offline MEPs of each block, there were significant differences (indicated by the brackets above the bars) between block 1 and blocks 2 and 3 (t_12_ = −3.928, −3.594; p = 0.006, 0.007; d = −1.089, −0.997) as well as a significant difference between blocks 2 and 3 (t_12_ = −2.466; p = 0.03; d = −0.684). N = 13.

Finally, the online and offline mean MEP amplitudes across each of the three tACS blocks were compared using a three-way rmANOVA. The FDI MEPs showed no main effects of BLOCK (F_2,24_ = 1.741, p = 0.197), STIMULATION (F_1,12_ = 2.681, p = 0.128), or any BLOCK × STIMULATION interactions (F_2,24_ = 0.125, p = 0.883). The ADM MEPs, on the other hand, showed a main effect of BLOCK (F = 11.394; p < 0.001; η^2^ = 0.073), with subsequent post-hoc t-tests confirming significant differences between block 1 and blocks 2 and 3 (t_12_ = − 3.928, −3.594; p = 0.006, 0.007; d = −1.089, −0.997) as well as a significant difference between blocks 2 and 3 (t_12_ = −2.466; p = 0.03; d = −0.684), supporting a gradual build-up in ADM MEP amplitude across blocks. However, similar to the FDI, there was no main effect of STIMULATION, although it was approaching significance (F_1,12_ = 4.287; p = 0.061), nor were there any BLOCK × STIMULATION interactions (F_2,24_ = 0.1, p = 0.905). There were no main effects of SESSION for either muscle (FDI: F_2,24_ = 1.558, p = 0.231; ADM: F_2,24_ = 1.443, p = 0.256); although there were significant BLOCK × SESSION interactions for both muscles (FDI: F = 3.075, p = 0.025, η^2^= 0.056; ADM: F = 3.269, p = 0.019, η^2^= 0.019). Post-hoc t-tests showed that the BLOCK × SESSION interaction for the ADM was driven by significant differences between blocks 1 and 3 for session 1 (t_12_ = −4.462, p = 0.001) and session 3 (t_12_ = − 4.779, p < 0.001), rather than a significant difference between sessions. However, the driving force for the FDI BLOCK × SESSION interaction could not be elucidated by the post-hoc t-tests.

### 3.3. Aftereffects of µ-tACS on µ/IMF EEG Power

The aftereffects of the tACS paradigm on both µ (8–13 Hz) and IMF (± 1 Hz) spectral power were assessed by comparing the pre-tACS, post-tACS, break, and echo period mean relative EEG powers using a two-way rmANOVA. As shown in Figure 5, there were significant main effects of TIME for both µ (F_6,66_ = 5.535, p < 0.001, η^2^ = 0.217) and IMF (F_6,66_ = 5.232, p < 0.001, η^2^ = 0.201), as well as significant TIME x SESSION interactions for both µ (F_12,132_ = 3.048, p < 0.001, η^2^ = 0.059) and IMF (F_2,22_ = 2.304, p = 0.011, η^2^ = 0.048); however, there were no main effects of SESSION for µ (F_2,22_ = 2.206, p = 0.134) or IMF (F_2,22_ = 2.309, p = 0.123). Post-hoc t-tests showed significant decreases in relative µ and IMF power from pre to echoes 1, 2 and 3 (µ: t_11_ = 9.172, 6.071, 5.322, p = <0.001, 0.002, 0.005, d = 2.648, 1.753, 1.536 respectively; IMF: t_11_ = 5.976, 5.779, 5.605, p = 0.002, 0.002, 0.003, d = 1.725, 1.668, 1.618 respectively) and a significant increase from break 2 to post (µ: t_11_ = −5.275, p = 0.005, d = −1.523; IMF: t_11_ = −5.205, p = 0.005, d = −1.503).

**Figure 5.**
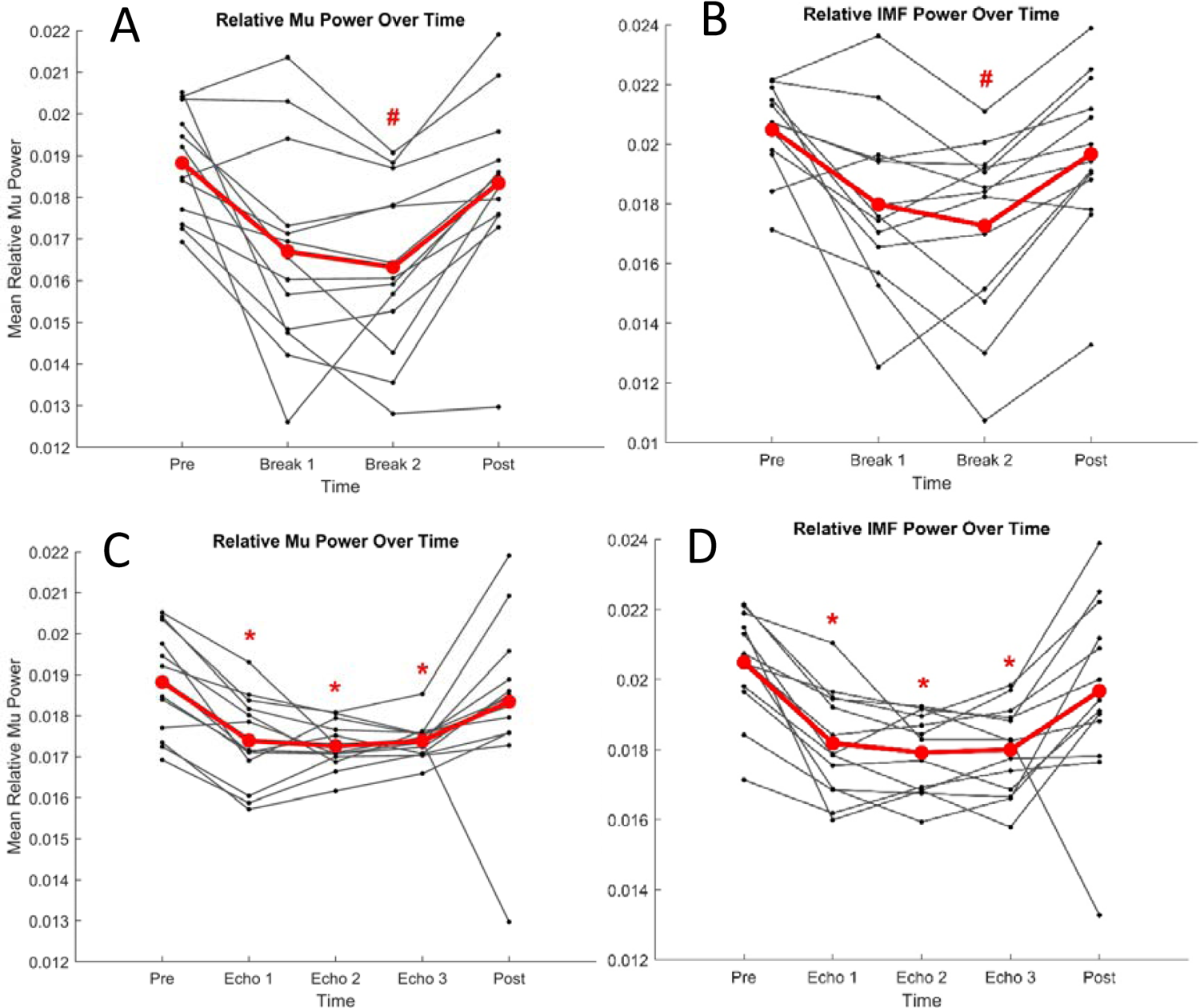
Individual Mean µ (8-13 Hz; A+C) and IMF (± 1 Hz; B+D) Relative Power for the Break (A+B) and Echo (C+D) Periods. Points represent each participant’s mean µ (8-13 Hz; A+C) and IMF (± 1 Hz; B+D) relative power from EEG activity acquired during the pre- and post-tACS (∼5 mins each) periods as well as the break (∼5 mins, A+B) and echo (∼2 mins; C+D) periods of each tACS block. Individual differences in µ/IMF relative power are represented by the solid lines. The global mean µ/IMF relative power for each period is represented by a red point, with the global mean difference between periods represented by a red line connecting the red points. There were significant decreases in relative µ and IMF power from pre to echoes 1, 2, and 3 (µ: t_11_ = 9.172, 6.071, 5.322, p = <0.001, 0.002, 0.005, d = 2.648, 1.753, 1.536 respectively; IMF: t_11_ = 5.976, 5.779, 5.605, p = 0.002, 0.002, 0.003, d = 1.725, 1.668, 1.618 respectively; *) as well as significant increases in µ and IMF power from break 2 to post (µ: t_11_ = −5.275, p = 0.005, d = −1.523; IMF: t_11_ = −5.205, p = 0.005, d = −1.503; #). N = 13.

### 3.4. Unaffected vs. Affected Sessions

We then performed each of the above analyses separately for the unaffected and affected sessions to assess the importance of precise IMF estimation for inducing entrainment and plastic aftereffects. The main findings of these separate analyses are summarised in Supplementary Table 1. Regarding the permutation analysis, the unaffected sessions remained significant at the group level for the ADM late online and echo MEPs (p = 0.0294 and 0.0035 respectively) and approached significance for the FDI early online and echo MEPs (p = 0.0605 and 0.0703 respectively), whilst the affected sessions were only significant for the FDI and ADM late online MEPs (p = 0.0215 and 0.0338 respectively) but were not significant for the echo MEPs of either muscle (p = 0.4261 and 0.1282 respectively). The pre-post increase in MEP amplitude remained significant for the unaffected and affected sessions for both the FDI (p = 0.018 and 0.007 for the unaffected and affected sessions respectively) and ADM (p = 0.006 and 0.002 respectively). In fact, the effect sizes were slightly greater for the affected sessions (d = −1.095 and −1.353 for the FDI and ADM respectively) compared to the unaffected sessions (d = −0.85 and −1.041 respectively). The only other change in terms of main effects was for the analysis comparing the offline MEP amplitudes of each block and the pre/post MEP amplitudes, where the affected sessions did not show a main effect of TIME for the FDI, although it approached significance (p = 0.085). The remaining analyses comparing MEP amplitudes and relative EEG powers across blocks showed no changes in main effects for the unaffected or affected sessions, although there were some changes to the significance of some of the post-hoc t-tests.

### 4. Discussion

Although animal studies (see review by Reato et al., 2013) have provided us with a cursory understanding of how tACS can shift the membrane potentials of pyramidal cells to alter their spike timing in a frequency- and phase-specific manner (Elyamany et al., 2021; Reato et al., 2010), it is less clear how these cellular effects translate into large-scale changes in network activity to induce some of the behavioural and cognitive effects of tACS seen in human studies (for reviews see, Antal & Paulus, 2013; Bland & Sale, 2019; Herrmann et al., 2016; Vosskuhl et al., 2018). Two primary mechanisms have been proposed thus far: entrainment of endogenous oscillatory activity to the frequency and phase of stimulation (Ali et al., 2013; Helfrich et al., 2014a,b; Huang et al., 2021; Witkowski et al., 2016) and neuroplastic aftereffects (Kasten et al., 2016; Neuling et al., 2013; Veniero et al., 2015; Vossen et al., 2015). The current study aimed to probe both these mechanisms in the context of µ-tACS using both phase-dependent TMS and EEG to assess the phasic (i.e., entrainment) effects on CSE and the sustained (i.e., neuroplastic) aftereffects on CSE and µ power. We report a sustained increase in both FDI and ADM MEP amplitudes as well as preliminary evidence for phasic entrainment of MEP amplitudes persisting briefly post-stimulation. We also report an acute decrease in relative µ/IMF EEG power, although there are some limitations to the experimental design that hinder the interpretation of these EEG results.

#### 4.1. Aftereffects of µ-tACS on CSE

Regarding the aftereffects of the µ-tACS paradigm on CSE, we observed significant increases in mean MEP amplitude from pre- to post-tACS for both the FDI (43.3% increase) and ADM (50% increase), in line with recent µ-tACS studies (Feurra et al., 2019; Fresnoza et al., 2018; Madsen et al., 2019a) and thus, supporting a facilitatory effect of µ oscillations on CSE (Bergmann et al., 2019; Karabanov et al., 2021; Ogata et al., 2019; Thies et al., 2018; Wischnewski et al., 2022). The response rate for this facilitatory effect was also relative;y consistent across participants, with 11 of the 13 participants demonstrating an increase in MEP amplitude (∼85% response rate), which represents, to our knowledge, the most effective means of increasing CSE across the multitude of plasticity-inducing paradigms currently in use (López-Alonso et al., 2014; Pellegrini et al., 2018; Veniero et al., 2015).

In fact, López-Alonso et al. (2014) assessed inter-individual variability with regards to the aftereffects of several commonly used plasticity-inducing paradigms (paired-associative stimulation, anodal transcranial direct current stimulation, and intermittent theta burst stimulation) on CSE across 56 participants and found that less than 50% of participants responded as expected to all 3 paradigms (39%, 45%, and 43% response rates respectively), akin to the response rates observed in previous studies (Hamada et al., 2012; Müller-Dahlhaus et al., 2008; Wiethoff et al., 2014). Furthermore, contrary to the present study, no significant changes in MEP amplitude were observed for any of the stimulation paradigms when the entire sample was analysed (i.e., both responders and non-responders). However, the response rate reported in the present study should also be interpreted with some caution due to the relatively small participant sample size.

High-frequency (i.e., > 1 Hz) repetitive TMS is also capable of plastically facilitating CSE (Chen and Seitz, 2001; Fitzgerald et al., 2006). However, similar to the other plasticity-inducing paradigms assessed by López-Alonso et al. (2014), it is also prone to significant inter- and intraindividual variability (Di Lazzaro et al., 2002; Maeda et al., 2000), with factors such as genetics (Hwang et al., 2015; Li Voti et al., 2011), menstrual cycle (Inghilleri et al., 2004), and even time of day (Sale et al., 2008) thought to contribute to this variability. Furthermore, tACS also holds a number of practical advantages over repetitive TMS, including: lower cost, greater portability, greater tolerance by subjects, and lower likelihood of seizure induction (Vosskuhl et al., 2018).

There were some slight differences in how these excitatory effects manifested for each muscle within the stimulation period. The FDI exhibited a relatively sharp increase in MEP amplitude from pre-tACS to block 1 that was not significantly different across the 3 blocks, whilst the ADM exhibited a more gradual increase in MEP amplitude that was not significantly different from pre-tACS until block 2 but was significantly different across blocks. This may suggest a potential ceiling effect for tACS-induced MEP amplitude facilitation. Again, however, these results should be interpreted with some caution due to the relatively small participant sample size, particularly with regards to the three-way repeated measures ANOVA comparing the online and offline MEP amplitudes across blocks.

It is of interest to note that these facilitatory effects are of a similar magnitude to the facilitatory effects observed in our most recent slow oscillatory tACS study (45.05% increase; Geffen et al., 2021) despite the differences in stimulation frequency, suggesting that these plastic aftereffects may not necessarily be frequency-dependent. When compared to β-tACS (15–25 Hz), which has been previously shown to facilitate resting CSE across multiple studies (Cancelli et al., 2015a; Cancelli et al., 2015b; Feurra et al., 2011; Feurra et al., 2013; Feurra et al., 2019; Heise et al., 2016), a recent meta-analysis by Wischnewski et al. (2019) reported that MEP amplitudes were only increased by β-tACS when using electrode montages with a more posterior location for the return electrode (e.g., M1-Pz or M1-Oz) but not when using a conventional M1-supraorbital region montage such as was used in the present study. This would seem to suggest differing effects for µ- and β-tACS depending on the exact stimulation site within the sensorimotor cortex or even across neighbouring regions (e.g., premotor cortex).

Because the present experiment did not include a negative control stimulation condition (e.g., sham stimulation), changes in MEP amplitude due to other factors (i.e., time and/or arousal effects) technically cannot be excluded. However, it seems highly unlikely that the increases in MEP amplitude reported here would be solely due to time and/or arousal effects, since a recent meta-analysis by Dissanayaka et al. (2018) found no significant effects of sham stimulation on MEP amplitude compared to baseline. Although only some of the studies assessed by this meta-analysis investigated tACS specifically, all of the studies used a comparable sham stimulation condition, and thus, they can all be used to make inferences about changes in MEP amplitude solely due to time and/or arousal. This therefore provides a compelling null comparator for the significant increase in MEP amplitude by µ-tACS that we report here.

#### 4.2. Phasic Effects of µ-tACS on CSE

Contrary to the plastic aftereffects of µ-tACS on CSE that were relatively consistent across participants, the phasic effects on CSE showed a greater degree of interindividual variability. Despite this however, 8 out of 13 participants exhibited significant phasic entrainment of FDI and/or ADM MEP amplitudes for at least one of the epochs. Furthermore, the group level analyses showed significant entrainment for the offline echo FDI and ADM MEPs as well as the late online ADM MEPs. To the best of our knowledge, this is the first evidence of phasic entrainment of MEP amplitudes by tACS persisting beyond stimulation, contrasting with the only other µ-tACS study that has explicitly assessed phasic entrainment of MEP amplitudes, which failed to show such an effect of phase on MEP amplitudes (Madsen et al., 2019). However, contrary to previous EEG experiments by Helfrich et al., (2014a,b), there was no correlation between the strength of entrainment and the magnitude of aftereffects, suggesting that these two effects are at least partially dissociable (Veniero et al., 2015; Vossen et al., 2015). There was also some notable intraindividual variability across the epochs, with only two participants exhibiting significant entrainment across two of the epochs (early and late online ADM MEPs) and no participants exhibiting entrainment across all epochs. It should be noted, however, that entrainment was not expected to occur for the early online MEPs due to the short duration of stimulation at this epoch (∼100–200 ms post-tACS onset).

The inter- and intraindividual variability observed for the permutation analysis is likely related to the inherent trial-to-trial variability of MEP amplitudes (Capaday, 2021), which can be impacted by several factors both within and across sessions (Darling et al., 2006). This includes physiological factors, such as fluctuations in resting brain activity (discussed further in the following section; Vidaurre et al., 2017; Zalesky et al., 2014) or unprovoked motor imagery (Gandevia and Rothwell, 1987; Hashimoto and Rothwell, 1999; Niyazov et al., 2005), as well as methodological factors, such as subtle changes in the position and/or orientation of the TMS coil (Grey and van de Ruit, 2017) due to the manual targeting of the M1 hotspot as opposed to more advanced targeting techniques using neuroimaging (e.g., the proprietary Neuronavigation system; Sparing et al., 2010; Thielscher et al., 2012). Although we initially attempted to include Neuronavigation in the pilot sessions of the present study, this proved challenging (and ultimately unachievable) due to the complexity of performing concurrent tACS, TMS, and EEG. However, it is worth noting that the trial-to-trial variability from these factors would have likely reduced the magnitude of any phasic effects of tACS on MEP amplitude (i.e., they are unlikely to produce a false positive), yet we still report evidence supporting entrainment of MEP amplitudes despite this.

#### 4.3. Aftereffects of µ-tACS on µ/IMF EEG Power

Regarding the aftereffects on EEG power, we observed significant decreases in relative µ and IMF power from pre-tACS to each of the echo periods, as well as significant increases in relative µ and IMF power from break 2 to post-tACS. Although this contradicts our hypothesised effects on EEG power, it is possible that these decreases in relative power for the echo periods reflect an acute homeostatic/refractory effect immediately following stimulation (Ketz et al., 2018). Alternatively, since tACS was applied using an open-loop paradigm as opposed to a closed-loop paradigm (i.e., stimulation was not synchronised to the endogenous phase), these decreases may reflect an initial desynchronisation of the endogenous µ rhythm due to a mismatch between the endogenous and exogenous (i.e., tACS) phases. This in turn may have gradually increase the susceptibility of the network to entrainment (i.e., resynchronisation) with respect to tACS phase, since desynchronisation of the targeted endogenous oscillation has been proposed to enhance entrainment to tACS phase (Krause et al., 2022; Lefebvre et al., 2017). Although this explanation is purely speculative, it may explain why we observed a decrease in µ power despite observing phasic entrainment. However, there are some limitations to the experimental design that hinder the interpretation of these EEG results.

The first of these limitations is the wakeful rest condition that was used in place of a behavioural test. Although this resting condition was initially chosen to avoid any external influence of a behavioural test on tACS-induced entrainment and/or plasticity, there is growing evidence suggesting that resting brain states can be highly variable both across and within individuals (Vidaurre et al., 2017; Zalesky et al., 2014), and thus, changes in EEG power solely due to time cannot be excluded. In fact, a recent functional magnetic resonance imaging (fMRI) study by Meer et al. (2020) characterised 10 distinct states of cortical activation and found that although resting-state brain dynamics are predominantly bistable between two of these states, the exact brain state dynamics varied significantly both across and within individuals. Importantly, a previous EEG study by Freyer et al. (2009) has suggested that this bistability in resting brain state dynamics reflects non-classic bursting between high- and low-amplitude periods of visual alpha power. Therefore, assuming that resting sensorimotor µ power fluctuates in a similar bistable fashion, it is possible that the wakeful rest condition used in the present study did not sufficiently control for these dynamic changes in endogenous µ power.

The dependency of tACS effects on endogenous brain states has been a topic of much interest for researchers in this field. It has recently been proposed that exogenous stimulation competes with endogenous oscillators for control of neuronal spike timing and that tACS-induced entrainment may be weakened if spike timing is already strongly entrained to the endogenous oscillators (Krause et al., 2022). Furthermore, a computational study by Lefebvre et al. (2017) has suggested that decreases in endogenous oscillatory power from certain behavioural tasks may enhance tACS-induced entrainment by increasing the range of the Arnold tongue compared to rest, allowing entrainment across a wider range of stimulation frequencies and increasing peak power when stimulating close to the resonant frequency. This inverse relationship between endogenous power and tACS effects has also been demonstrated for the plastic aftereffects of tACS, with human studies reporting significant tACS aftereffects when endogenous power is low (Geffen et al., 2021; Neuling et al., 2013) but not when endogenous power is high (Neuling et al., 2013).

Furthermore, a concurrent tACS–fMRI study by Vosskuhl et al. (2016) found that alpha-tACS may reduce task-induced blood oxygenation level dependent (BOLD) responses to visual targets but does not seem to modulate the resting state BOLD signal. This suggests that tACS alone cannot induce a BOLD response de novo but it may still be able to induce changes in BOLD activity by modulating an existing BOLD response, in the same way that low-intensity tACS is unlikely to induce a neural oscillation that is not already present in the cortex but can modulate the power of endogenous oscillations that are prominent in the network (Ali et al., 2013; Ketz et al., 2018; Liu et al., 2018).

The second limitation hindering the interpretation of the EEG results was the EEG electrode montage used to estimate µ power. A recent study by Karabanov et al. (2021) has suggested that EEG electrode montage may play a crucial role in detecting relationships between µ power and MEP amplitude, reporting a positive correlation between µ power and MEP amplitude when using a Laplacian montage centred around M1 (Thies et al., 2018) that was not present when using radial source projection. Importantly, this Laplacian montage on average has a more posterior location (i.e., closer to S1) compared to the source projection (i.e., closer to M1), supporting the notion that the positive relationship between µ power and MEP amplitude reflects an increase in µ activity in S1 that results in disinhibition of feed-forward inhibitory inputs from S1 to M1 (Bergmann et al., 2019). Unfortunately, however, it was not possible to utilise this Laplacian montage in the present study due to the placement of the tACS pad over M1, and thus, our estimation of endogenous µ power may not have been optimised. Therefore, given that µ power was not able to be assessed with a Laplacian montage and that dynamic changes in resting brain states may not have been adequately controlled for, the changes in relative µ power observed here should be interpreted cautiously.

#### 4.4. Impact of Precise IMF estimation

Because of the errors in IMF estimation that only became apparent post-data collection, some of the sessions had a mismatch between the stimulation frequency (i.e., the initial IMF estimate) and the endogenous eigenfrequency (i.e., the revised IMF estimate). Therefore, we chose to perform separate analyses for the sessions that were affected by these mismatches and the sessions that were unaffected so that we could assess the importance of precise IMF estimation for inducing entrainment and plastic aftereffects. However, it should be noted that any differences in significance (or lack thereof) between the complete analysis and these separate analyses could simply be due to the reduced number of analysed sessions combined with the fact that those sessions had to be averaged for each participant.

The fact that the affected sessions did not show significant entrainment echoes at the group level for either muscle (although they showed significant late online entrainment) suggests that precise IMF estimation may increase the likelihood of inducing entrainment echoes using tACS, in line with the proposed resonance dynamics (i.e., the Arnold tongue) for tACS-induced entrainment (Ali et al., 2013; Huang et al., 2021; Liu et al., 2018; Schutter and Wischnewski, 2016; Thut et al., 2017; Vosskuhl et al., 2018). However, the plastic facilitation of MEP amplitudes did not seem to be significantly impacted by the precision of IMF targeting, in-line with previous tACS studies demonstrating similar plastic aftereffects at stimulation frequencies outside the µ frequency band (8–13 Hz), such as slow-wave (0.75 Hz; Geffen et al., 2021) and β-tACS (15–25 Hz; Wischnewski et al., 2019). In fact, the affected sessions even showed a slightly greater facilitation of MEP amplitudes compared to the unaffected sessions. These findings further support the notion that entrainment and plastic aftereffects are at least partially dissociable (Veniero et al., 2015; Vossen et al., 2015). The changes in relative µ/IMF power were mostly unaffected by the precision of IMF targeting, with no differences in main effects for the power analyses between the unaffected and affected sessions. Again, however, these EEG results should be interpreted cautiously because of the limitations discussed in the previous section.

## 5. Conclusion

To summarise, we present preliminary evidence supporting phasic entrainment of MEP amplitudes persisting beyond stimulation and have also replicated the sustained facilitation of MEP amplitudes observed in previous µ-tACS studies. These findings have important implications in the research and clinical domains as it demonstrates that tACS can effectively modulate neural activity by entraining CSE to match the frequency and phase of stimulation as well as inducing plastic aftereffects on CSE, thus supporting the two primary mechanisms proposed to underly the behavioural effects of tACS. However, the inter- and intraindividual variability observed for the entrainment effects and the changes in relative EEG power warrants further experimentation with improved IMF estimation and a suitable behavioural task to control for differences in resting brain states.

## Acknowledgements

We would like to thank Marc Mosimann for his assistance with setting up the Biosemi ActiveTwo EEG hardware and software.

## Abbreviations

tACS: Transcranial Alternating Current Stimulation

TMS: Transcranial Magnetic Stimulation

CSE: Corticospinal Excitability

IMF: Individual µ (Mu) Frequency

EEG: Electroencephalography

MEG: Magnetoencephalography

EMG: Electromyography

MEP: Motor Evoked Potential

M1_HAND_: Hand Area of the Primary Motor Cortex

S1: Primary Somatosensory Cortex

FDI: first dorsal interosseous

ADM: abductor digiti minimi

fMRI: Functional Magnetic Resonance Imaging

BOLD: Blood Oxygenation Level Dependent

## Supplementary Material

**Supplementary Figure 1.**
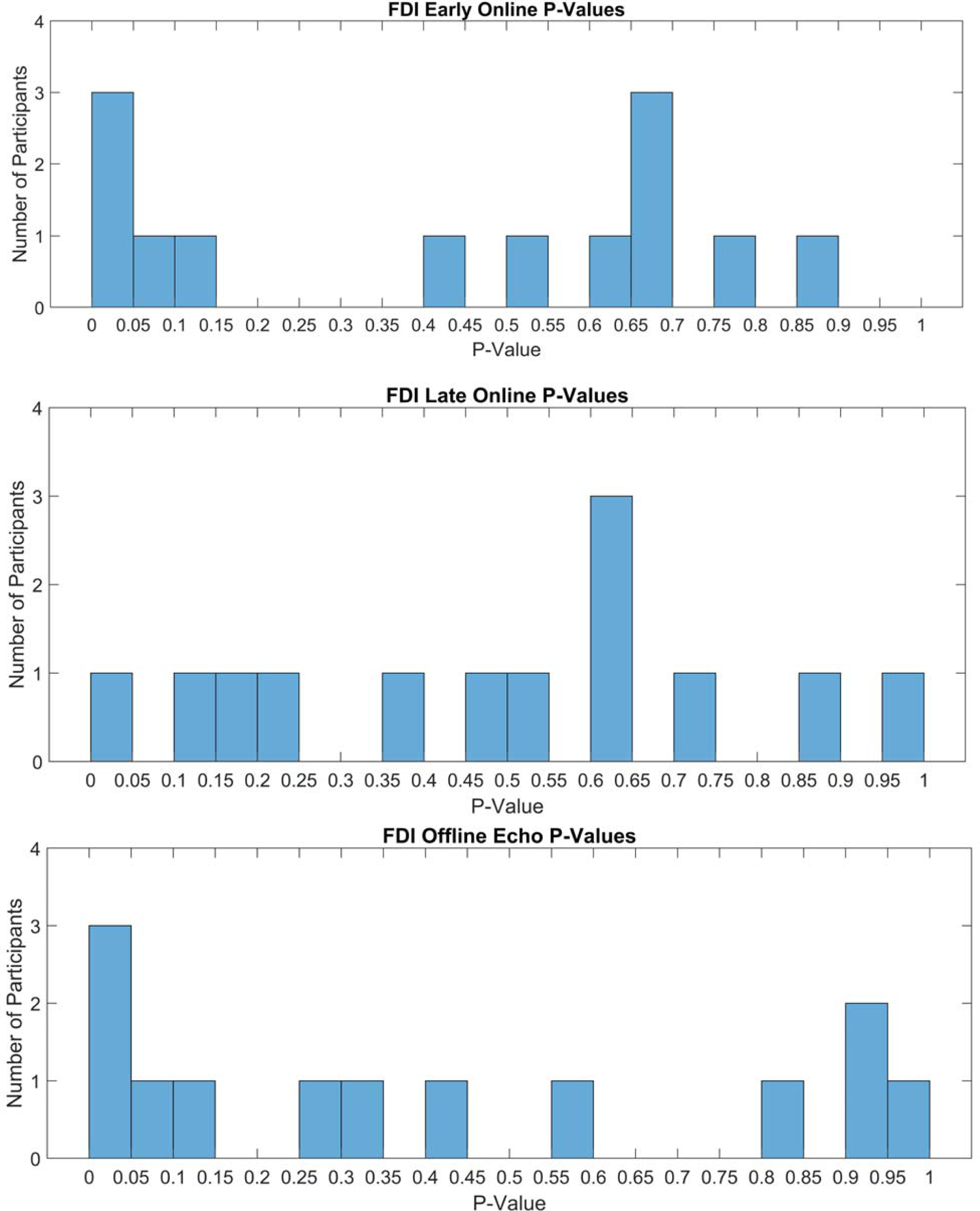
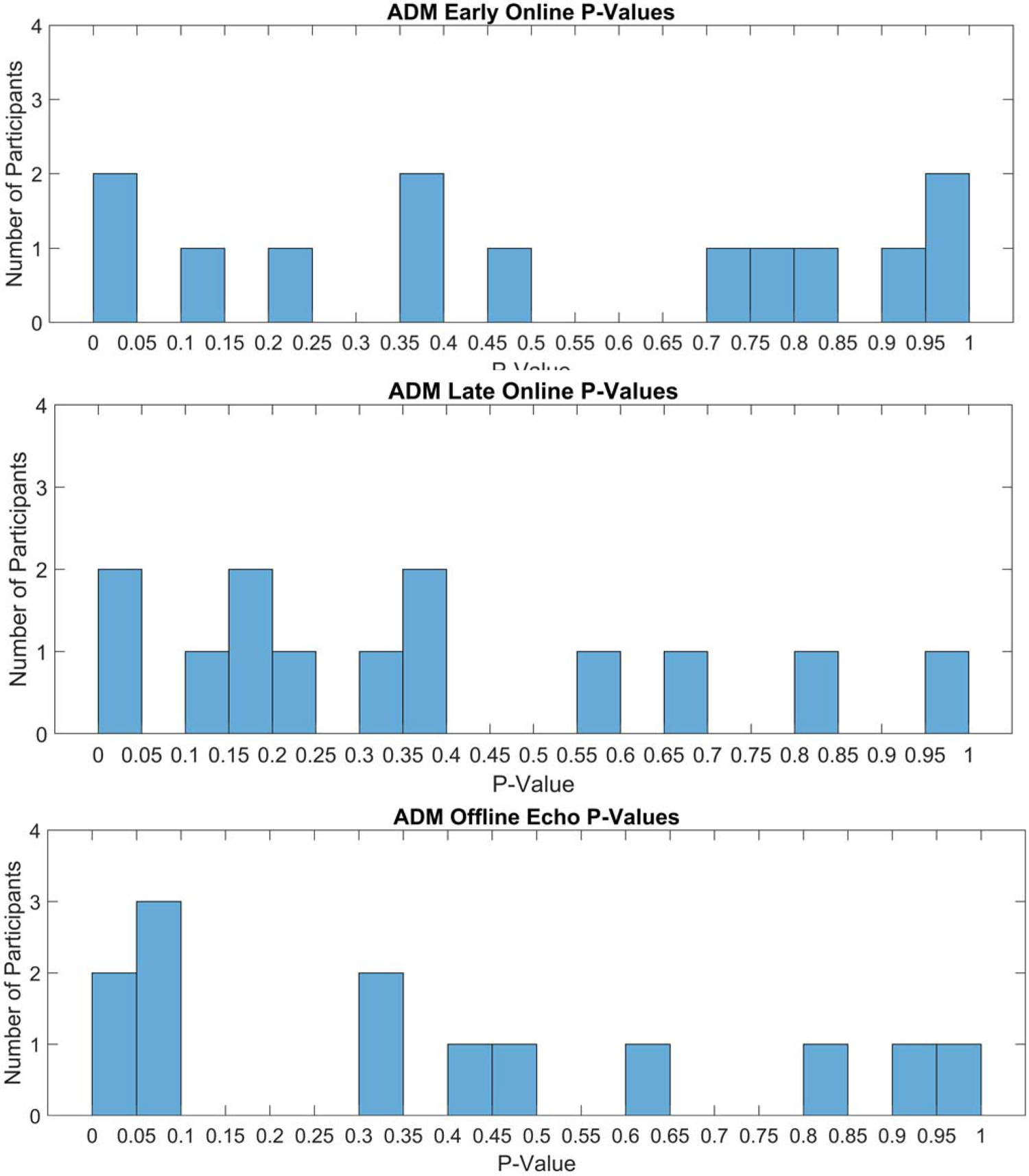
Histograms of Each Participant’s Permutation Analysis P-Values for the FDI (Top) and ADM (Bottom) MEPs. Bars represent the number of participants with P-values within the range specified on the x-axis for early online (A/D), late online (B/E), and offline echo (C/F) FDI/ADM MEPs.

---

**Supplementary Table 1.**
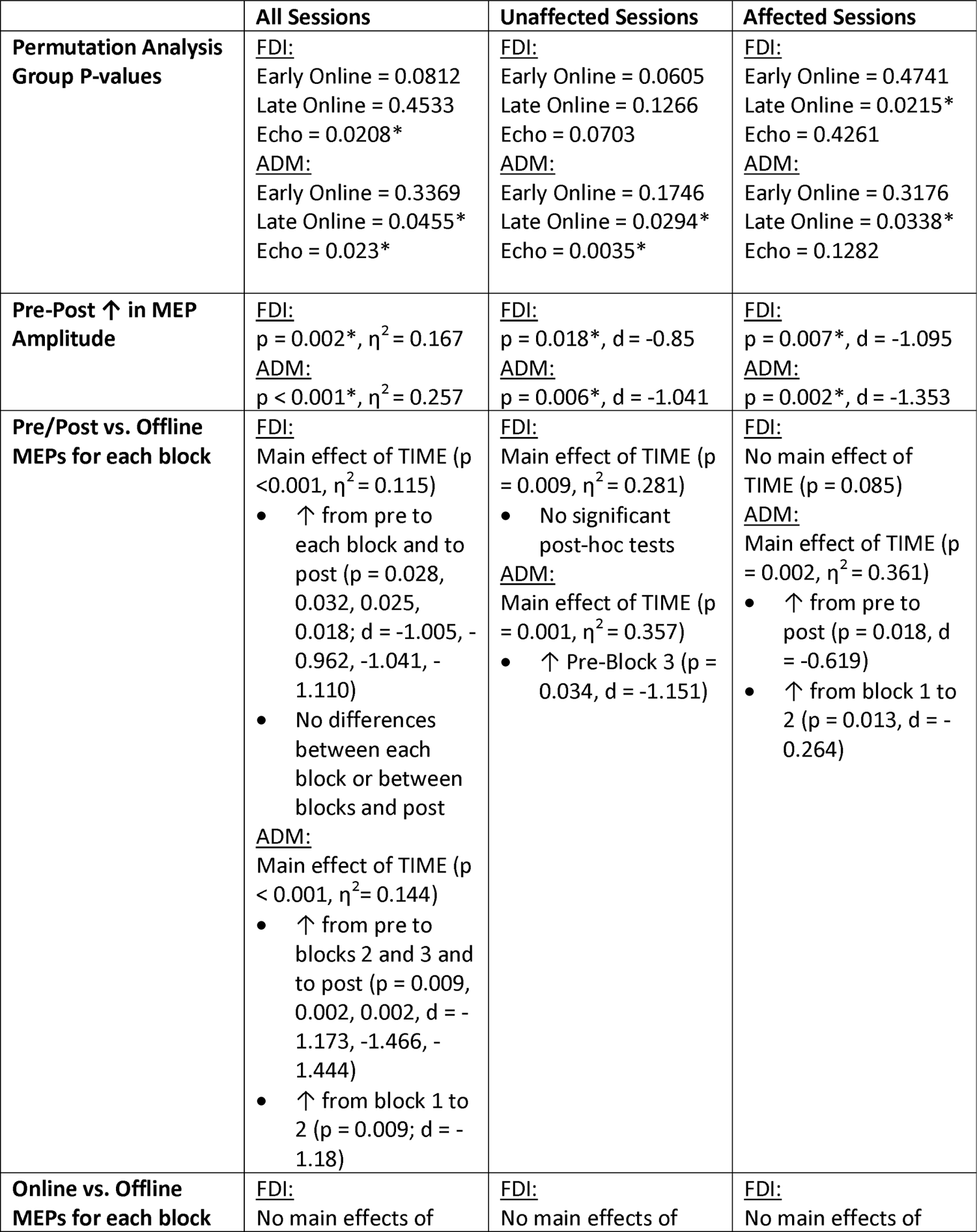
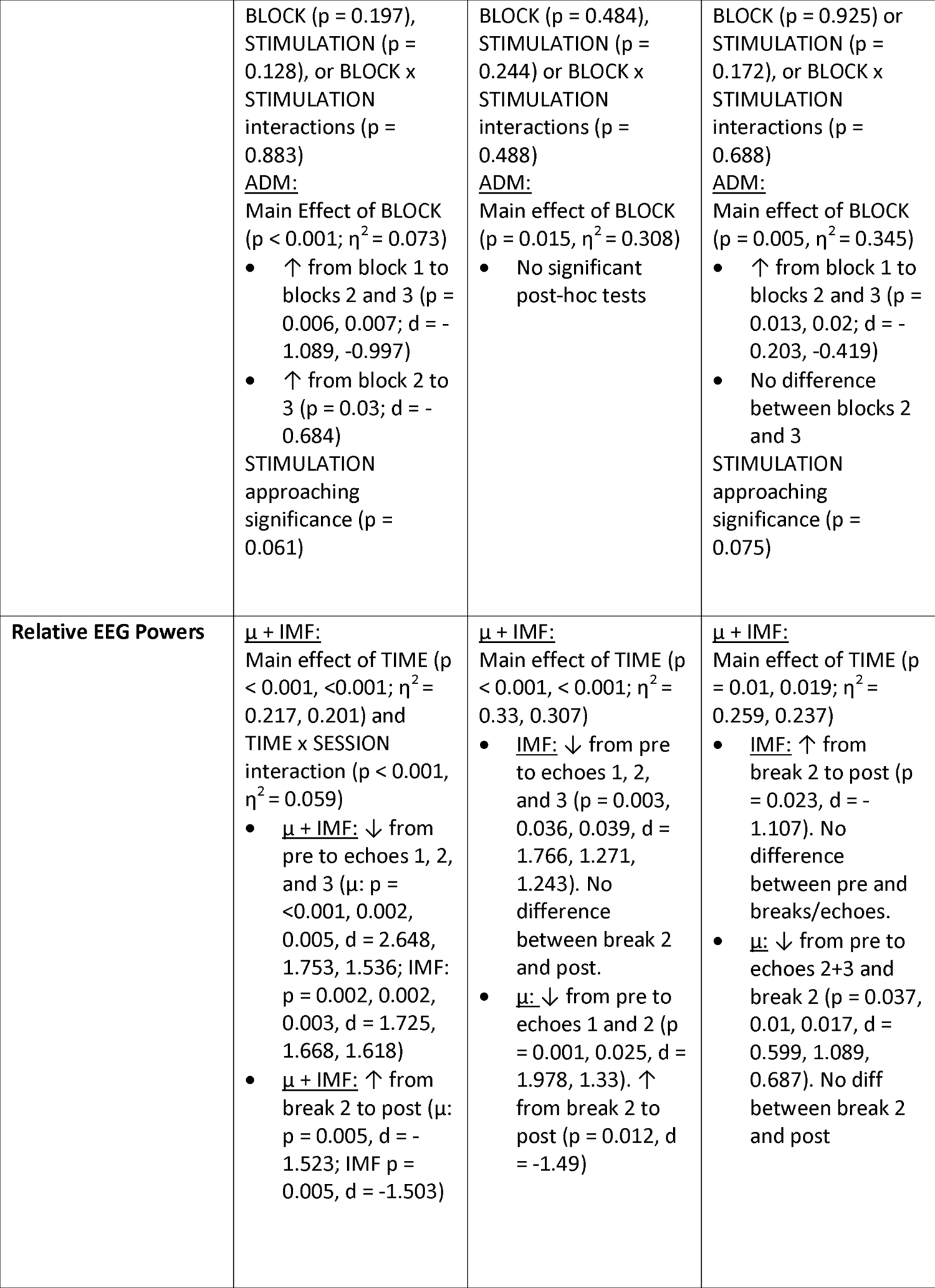
Summary of Results for All Sessions, Sessions with no Difference Between Initial and Revised IMF Estimates (Unaffected Sessions), and Sessions with a Difference Between Initial and Revised IMF Estimates (Affected Sessions). Whilst the affected sessions showed significant entrainment for the late online MEPs (p = 0.0215 and 0.0338 for FDI and ADM respectively), they did not show significant entrainment echoes for either muscle. There was no change in significance for the affected sessions with regards to the pre-post increase in MEP amplitude, although there was a loss of main effect of TIME for the analysis comparing the pre/post MEP amplitudes against the offline MEPs of each tACS block. The remaining analyses showed no changes to the main effects for the affected sessions, although there were some changes to the significance of some of the post-hoc t-tests.

