## Supplementary figures and images for "Mu-Transcranial Alternating Current Stimulation Induces Phasic Entrainment and Plastic Facilitation of Corticospinal Excitability"

### Supplementary Figure 1A

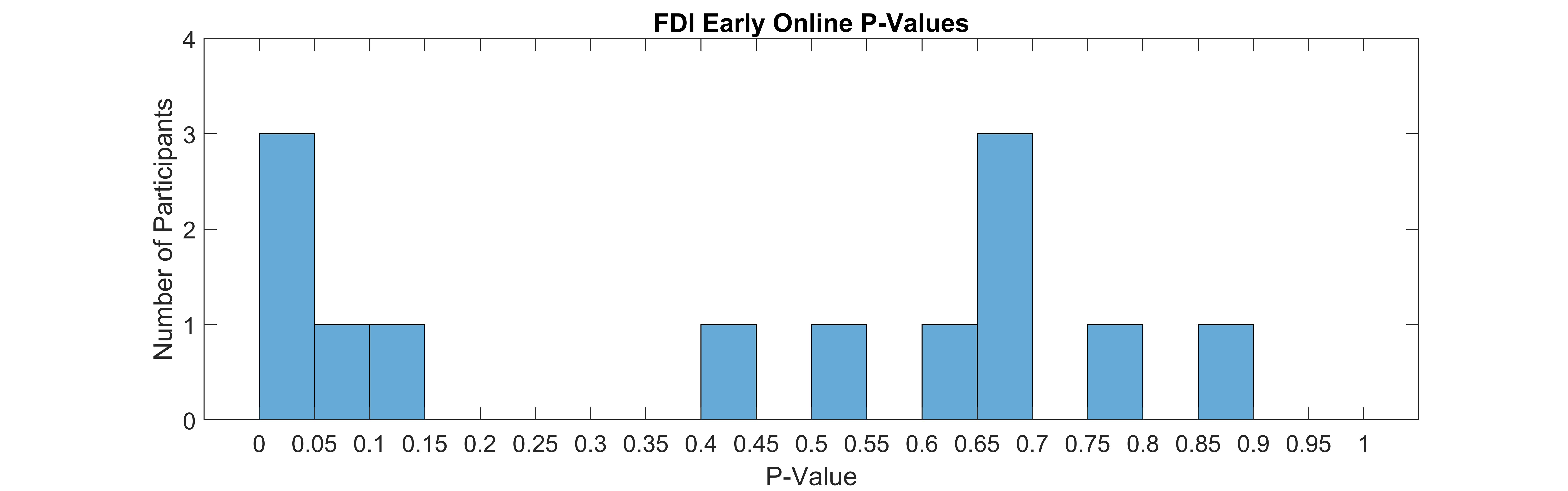

### Supplementary Figure 1B

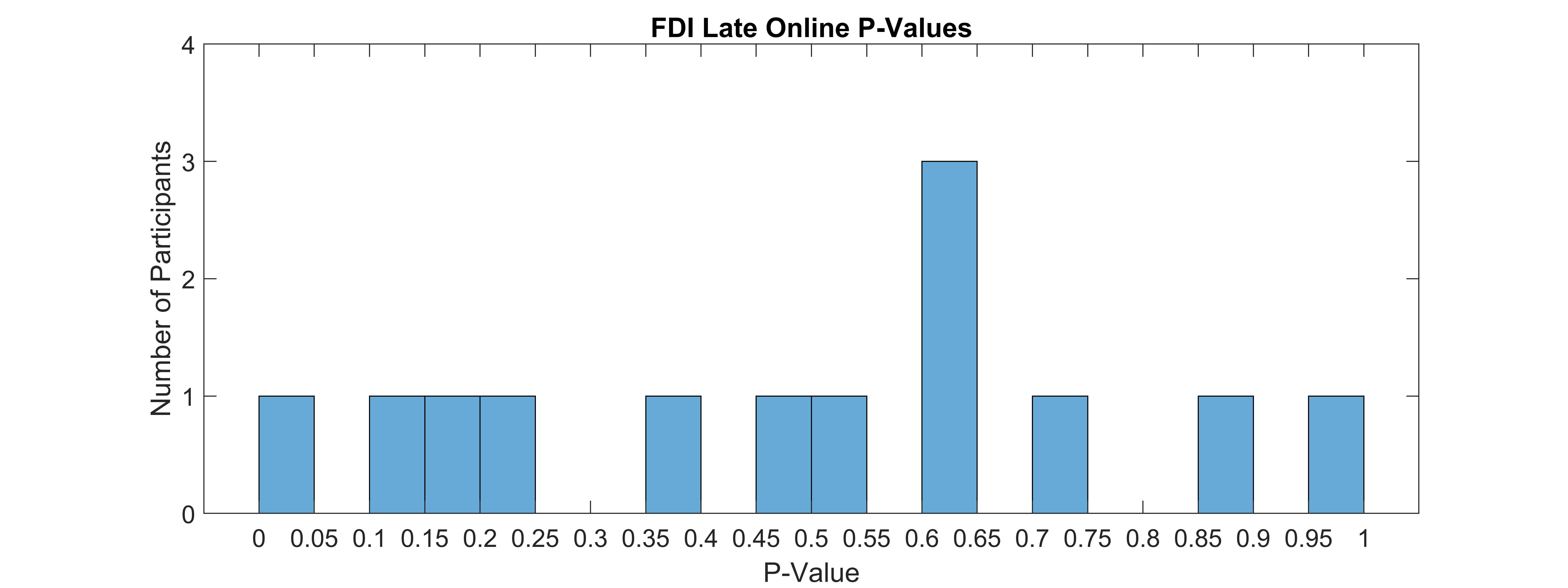

### Supplementary Figure 1C

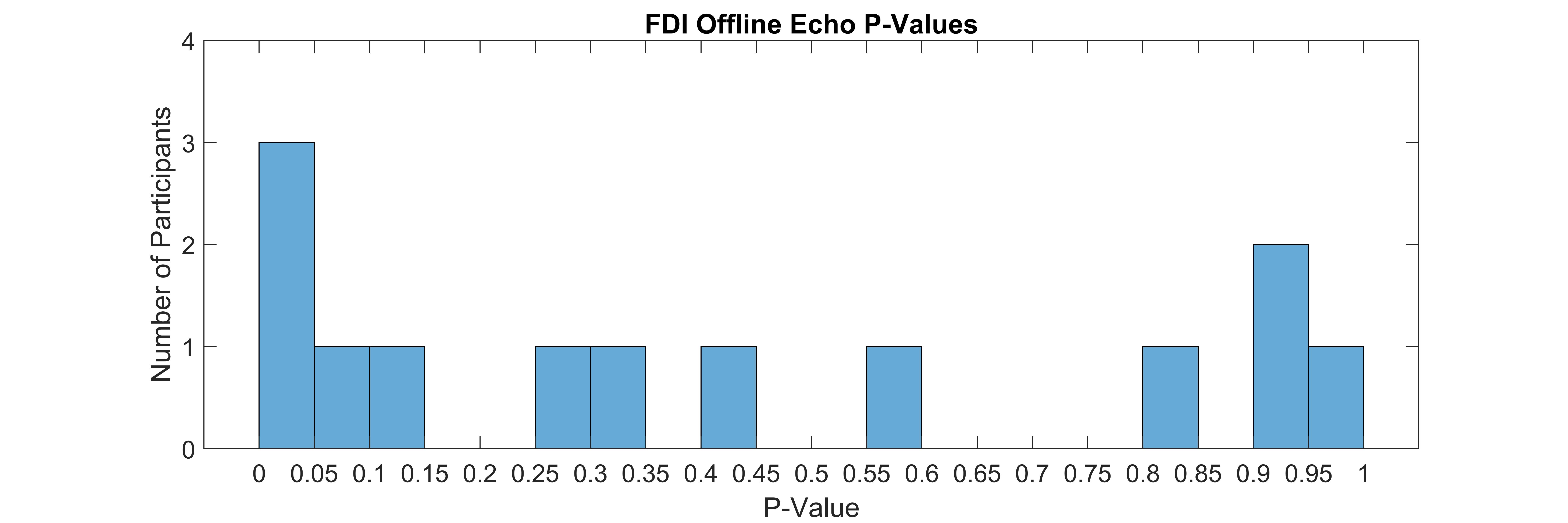

### Supplementary Figure 1D

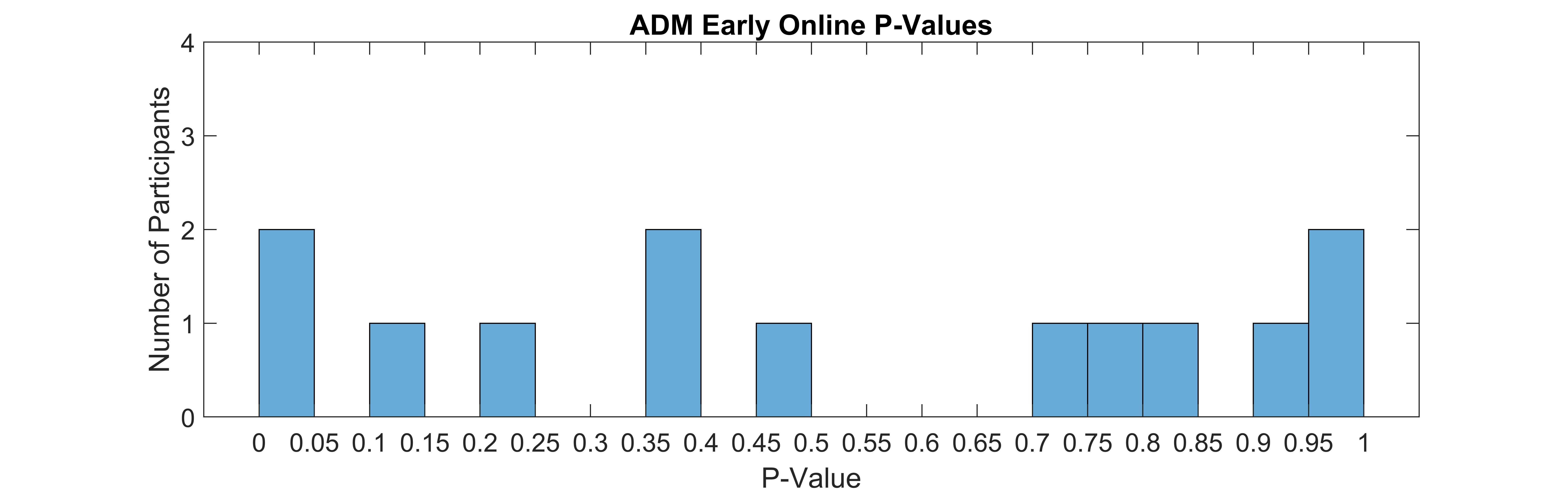

### Supplementary Figure 1E

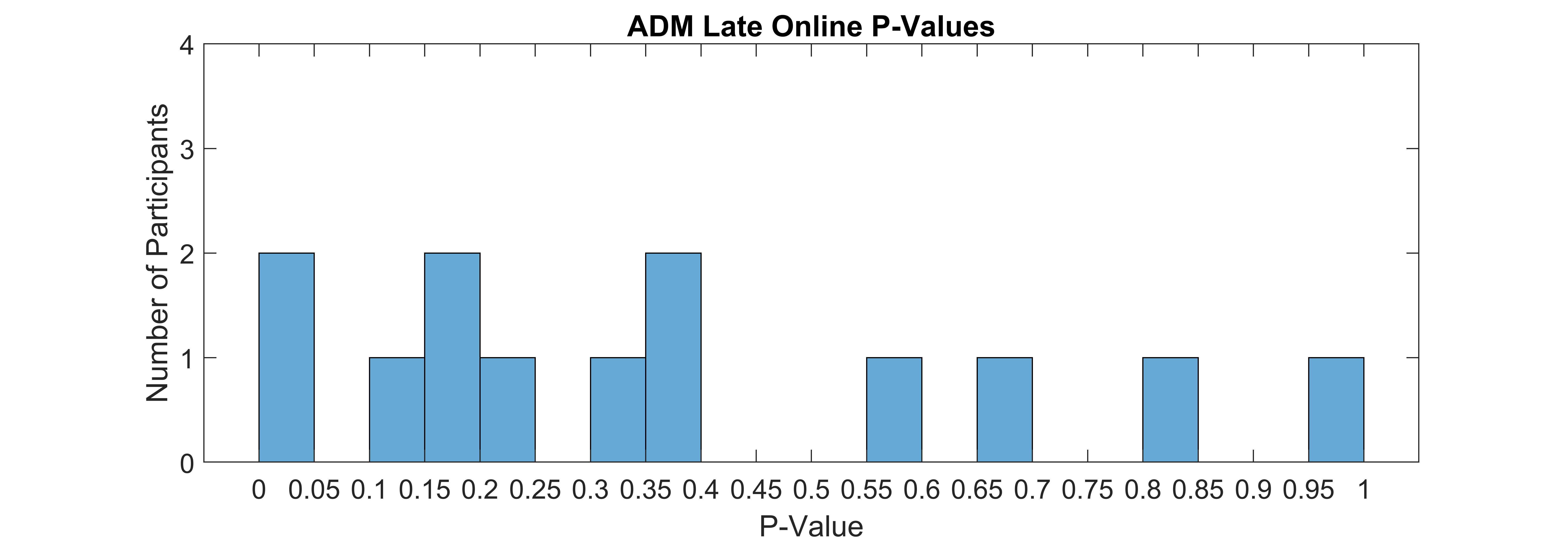

### Supplementary Figure 1F

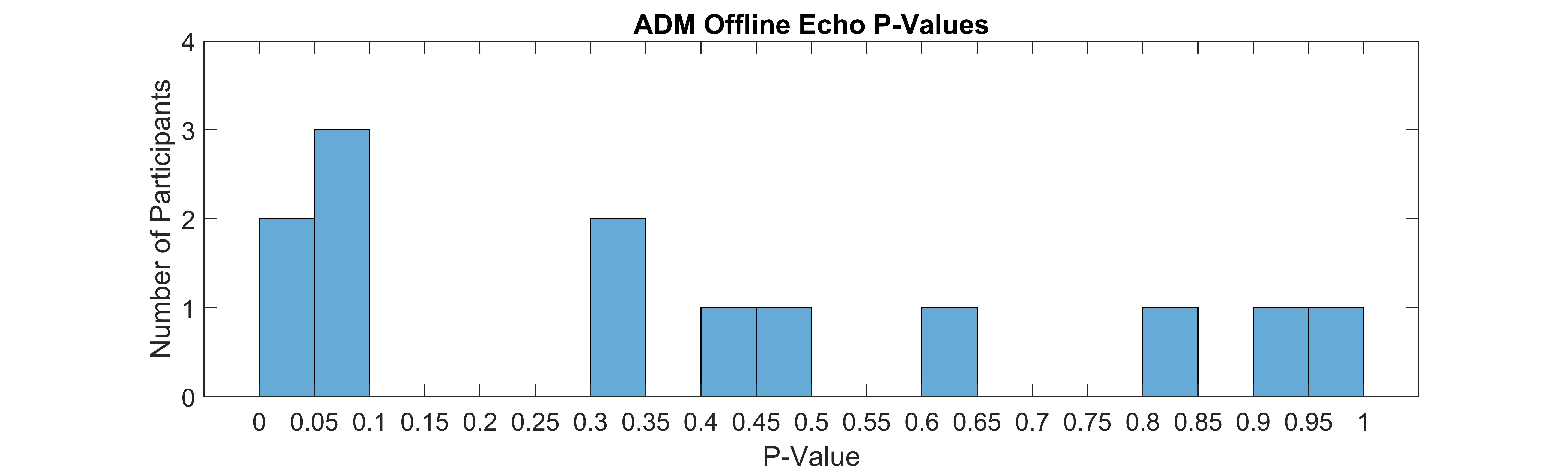
