## Supplementary Table 1 for "Mu-Transcranial Alternating Current Stimulation Induces Phasic Entrainment and Plastic Facilitation of Corticospinal Excitability"

**Supplementary Table 1.** **Summary of Results for All Sessions, Sessions with no Difference Between Initial and Revised IMF Estimates (Unaffected Sessions), and Sessions with a Difference Between Initial and Revised IMF Estimates (Affected Sessions).** Whilst the affected sessions showed significant entrainment for the late online MEPs (p = 0.0215 and 0.0338 for FDI and ADM respectively), they did not show significant entrainment echoes for either muscle. There was no change in significance for the affected sessions with regards to the pre-post increase in MEP amplitude, although there was a loss of main effect of TIME for the analysis comparing the pre/post MEP amplitudes against the offline MEPs of each tACS block. The remaining analyses showed no changes to the main effects for the affected sessions, although there were some changes to the significance of some of the post-hoc t-tests.

|  | **All Sessions** | **Unaffected Sessions** | **Affected Sessions** |
| --- | --- | --- | --- |
| **Permutation Analysis Group P-values** | FDI:  Early Online = 0.0812  Late Online = 0.4533  Echo = 0.0208*  ADM:  Early Online = 0.3369  Late Online = 0.0455*  Echo = 0.023* | FDI:  Early Online = 0.0605  Late Online = 0.1266  Echo = 0.0703  ADM:  Early Online = 0.1746  Late Online = 0.0294*  Echo = 0.0035* | FDI:  Early Online = 0.4741  Late Online = 0.0215*  Echo = 0.4261  ADM:  Early Online = 0.3176  Late Online = 0.0338*  Echo = 0.1282 |
| **Pre-Post ↑ in MEP Amplitude** | FDI:  p = 0.002*, η^2^ = 0.167  ADM:  p < 0.001*, η^2^ = 0.257 | FDI:  p = 0.018*, d = -0.85  ADM:  p = 0.006*, d = -1.041 | FDI:  p = 0.007*, d = -1.095  ADM:  p = 0.002*, d = -1.353 |
| **Pre/Post vs. Offline MEPs for each block** | FDI:  Main effect of TIME (p <0.001, η^2^ = 0.115)   - ↑ from pre to each block and to post (p = 0.028, 0.032, 0.025, 0.018; d = -1.005, -0.962, -1.041, -1.110) - No differences between each block or between blocks and post   ADM:  Main effect of TIME (p < 0.001, η^2^= 0.144)   - ↑ from pre to blocks 2 and 3 and to post (p = 0.009, 0.002, 0.002, d = -1.173, -1.466, - 1.444) - ↑ from block 1 to 2 (p = 0.009; d = -1.18) | FDI:  Main effect of TIME (p = 0.009, η^2^ = 0.281)   - No significant post-hoc tests   ADM:  Main effect of TIME (p = 0.001, η^2^ = 0.357)   - ↑ Pre-Block 3 (p = 0.034, d = -1.151) | FDI:  No main effect of TIME (p = 0.085)  ADM:  Main effect of TIME (p = 0.002, η^2^ = 0.361)   - ↑ from pre to post (p = 0.018, d = -0.619) - ↑ from block 1 to 2 (p = 0.013, d = -0.264) |
| **Online vs. Offline MEPs for each block** | FDI:  No main effects of BLOCK (p = 0.197), STIMULATION (p = 0.128), or BLOCK x STIMULATION interactions (p = 0.883)  ADM:  Main Effect of BLOCK (p < 0.001; η^2^ = 0.073)   - ↑ from block 1 to blocks 2 and 3 (p = 0.006, 0.007; d = -1.089, -0.997) - ↑ from block 2 to 3 (p = 0.03; d = -0.684)   STIMULATION approaching significance (p = 0.061) | FDI:  No main effects of BLOCK (p = 0.484), STIMULATION (p = 0.244) or BLOCK x STIMULATION interactions (p = 0.488)  ADM:  Main effect of BLOCK (p = 0.015, η^2^ = 0.308)   - No significant post-hoc tests | FDI:  No main effects of BLOCK (p = 0.925) or STIMULATION (p = 0.172), or BLOCK x STIMULATION interactions (p = 0.688)  ADM:  Main effect of BLOCK (p = 0.005, η^2^ = 0.345)   - ↑ from block 1 to blocks 2 and 3 (p = 0.013, 0.02; d = -0.203, -0.419) - No difference between blocks 2 and 3   STIMULATION approaching significance (p = 0.075) |
| **Relative EEG Powers** | µ + IMF:  Main effect of TIME (p < 0.001, <0.001; η^2^ = 0.217, 0.201) and TIME x SESSION interaction (p < 0.001, η^2^ = 0.059)   - µ + IMF: ↓ from pre to echoes 1, 2, and 3 (µ: p = <0.001, 0.002, 0.005, d = 2.648, 1.753, 1.536; IMF: p = 0.002, 0.002, 0.003, d = 1.725, 1.668, 1.618) - µ + IMF: ↑ from break 2 to post (µ: p = 0.005, d = -1.523; IMF p = 0.005, d = -1.503) | µ + IMF:  Main effect of TIME (p < 0.001, < 0.001; η^2^ = 0.33, 0.307)   - IMF: ↓ from pre to echoes 1, 2, and 3 (p = 0.003, 0.036, 0.039, d = 1.766, 1.271, 1.243). No difference between break 2 and post. - µ: ↓ from pre to echoes 1 and 2 (p = 0.001, 0.025, d = 1.978, 1.33). ↑ from break 2 to post (p = 0.012, d = -1.49) | µ + IMF:  Main effect of TIME (p = 0.01, 0.019; η^2^ = 0.259, 0.237)   - IMF: ↑ from break 2 to post (p = 0.023, d = -1.107). No difference between pre and breaks/echoes. - µ: ↓ from pre to echoes 2+3 and break 2 (p = 0.037, 0.01, 0.017, d = 0.599, 1.089, 0.687). No diff between break 2 and post |
